# A hierarchy of spatial confinements governs chromatin dynamics and genomic encounters

**DOI:** 10.64898/2026.07.12.738087

**Authors:** Megan Aubrey, Arpita Prusy, Olga K. Dudko, Cornelis Murre

## Abstract

Antibody diversity is generated through stochastic rearrangement of the immunoglobulin heavy chain locus (Igh) involving the recombination of variable (V_H_), diversity (D_H_) and joining (J_H_) gene segments. How coding elements within the Igh locus locate one another in the complex nuclear environment is not understood. Here, we sought to identify the molecular mechanisms and physical principles that govern V_H-_D_H_J_H_ genomic encounters. We found that transcription imposed a local confinement that stabilized interactions between spatially proximal genomic elements. The loop anchor CTCF modestly constrained population-average chromatin motion, whereas cohesin-mediated loops established large-scale confinement and reinforced self-similarity of V_H-_D_H_J_H_ motion across spatial and temporal scales. Quantitative scaling arguments for first-passage times revealed that encounter frequencies between remote V_H-_D_H_J_H_ genomic regions are governed by the interplay of diffusivity and spatial proximity. Together, these findings show that the hierarchy of confinements imposed by transcription and loop extrusion provides the balance between stability and mobility required to regulate genomic encounter frequencies.

---

The mammalian genome is folded into well-defined hierarchical structures. On the scale of the nucleus, chromosomes segregate into chromosome territories ^1–3^. Individual chromosomes, in turn, cluster into transcriptionally-active (A) and transcriptionally-inactive (B) compartments ^3^. At a finer scale, the chromatin fiber is structured as topologically associated domains (TADs), or stacked rosettes of chromatin loops that are anchored, at least in part, by the architectural protein CTCF ^3–6^. The assembly of loops is mediated by cohesin, a protein complex that orchestrates loop extrusion^5,7–10^. Specifically, a component of the cohesin complex, named RAD21, is sequestered and activated across the chromatin landscape by NIPBL to initiate and maintain loop extrusion^11,12^. Once loaded, cohesin progressively extrudes chromatin loops until a pair of convergent CTCF sites is reached ^5^. Loop extrusion is halted by WAPL-mediated release of cohesin from the chromatin landscape ^13^. Loop domains within TADs often assemble into phase-separated transcriptional condensates, composed of transcription factors, non-coding transcripts, chromatin remodelers and co-activators ^14–17^. Recent studies have explored the role of cohesin in modulating chromatin dynamics ^18,19^. These studies revealed that chromatin loops are dynamic structures, with most loops existing in a partially extruded state rather than bridging two convergent CTCF binding elements ^19^. RAD21-depletion and, to a smaller degree, CTCF-depletion increased spatial distances and the mean squared displacement of paired CTCF sites separated by a 500 kbp genomic region spanning a singular loop domain ^19^. Conversely, WAPL-depletion resulted in a decrease in spatial distances and the mean squared displacement ^19^.

B cell antigen receptors are encoded by gene segments distributed across three distinct genomic regions including (i) the immunoglobulin heavy chain (Igh) locus, (ii) the immunoglobulin kappa (Igκ) locus and (iii) the immunoglobulin lambda (Igλ) locus. The heavy chain is encoded by the Igh locus, whereas the light chain is encoded by either the Igκ or Igλ locus. The Igh locus is comprised of variable (V), diversity (D), joining (J) and constant (C) regions. The Igκ and Igλ are comprised of V, J and C regions but lack D elements. Igh and Igλ locus recombination occurs by deletion whereas Vκ-Jκ rearrangement occurs either by deletion or inversion. The Igh and Igκ loci utilize different mechanisms to generate antibody diversity. Igh locus rearrangement is instructed by RAG scanning involving the cohesin machinery that pulls D_H_J_H_ elements along the chromatin until they align with V_H_ gene segments to undergo V_H-_D_H_J_H_ locus rearrangement and, to a smaller degree, diffusion of V_H_, D_H_ and J_H_ elements ^20,21^. Chromatin architecture plays a critical role in orchestrating V_H-_D_H_J_H_ rearrangement. Specifically, CTCF-bound sites located between the V_H_ and D_H_J_H_ elements sequester the distal V_H_ genes away from the D_H_J_H_ region to suppress aberrant V_HDH g_ene rearrangement ^22,23^ (Figure S1a) In developing pro-B cells, cohesin assembles a wide spectrum of long-range loops, which are anchored by hundreds of CTCF-bound sites that, in turn, are positioned in convergent orientation with CTCF-bound sites located at the Igh locus super-anchor (Figure S1a) ^23,24^. During the course of B cell progression, a decline in WAPL expression results in prolonged loop extension, yielding large loops, large-scale locus remodeling, and locus contraction that is closely associated with increased distal V_H-_D_H_J_H_ rearrangements ^21,25^. As a first step towards unraveling how V_H-_D_H_J_H_ dynamics is modulated during B cell development, we have developed strategies to track paired remote genomic interactions in live B-lineage cells ^26,27^. Analysis of the mean squared displacements and velocity autocorrelations of the genomic trajectories, in conjunction with Brownian-and Molecular Dynamics simulations of polymer models of the locus, revealed that V_H-_D_H_J_H_ motion is subdiffusive as the result of the imposed network of crosslinked chromatin chains, and that the chromatin network is poised near the boundary between gel and sol phases ^27^. Subdiffusive motion is characterized by mean squared displacement scaling slower than linearly with time, MSD ∼ Dτ^α^, where D is the diffusion coefficient describing the overall magnitude of chromatin motion and α < 1 is the anomalous scaling exponent characterizing subdiffusive chromatin motion arising from polymer connectivity and interactions in the chromatin environment.

Here we sought to determine how the physical environment of the Igh locus dictates subdiffusive V_H-_D_H_J_H_ dynamics. We found that cohesin-mediated chromatin loops imposed confinement at a global scale to (i) bring paired V_H_ and D_H_J_H_ elements into spatial proximity, (ii) constrain average V_H_-D_H_J_H_ motion, and (iii) reinforce similar patterns of V_H_-D_H_J_H_ motion across spatial and temporal scales. We found that transcription induced local-scale confinement to enrich for V_H_-D_H_J_H_ interactions involving V_H_ and D_H_J_H_ elements that were spatially co-localized through the global confinement of the loops. Consistent with previous observations, we found that CTCF occupancy modestly tightened the spatial constraint. We found that depletion of RAD21 perturbed V_H_-D_H_J_H_ chromatin mobility and Igh locus spatial confinement. Finally, we showed that V_H_-D_H_J_H_ chromatin diffusivity and spatial proximity were key factors that determine V_H_-D_H_J_H_ mean first-passage times. Taken together, our findings show that transcription and loop extrusion act cooperatively to assemble a hierarchical chromatin architecture that balances chromatin stability with dynamic genomic exploration to regulate V_H_-D_H_J_H_ encounter frequencies.

## RESULTS

### Non-coding transcription confines chromatin motion at the level of individual paired V_H_-D_H_J_H_ elements

Numerous studies have demonstrated that V_H_-D_H_-J_H_ rearrangement is closely associated with non-coding transcription ^28–34^. Prior to rearrangement, non-coding transcription is initiated across the D_H_-J_H_ region and the Eμ intronic enhancer ^31–33^. Following D_H_-J_H_ rearrangement, sense-and antisense-transcription is initiated in the genomic regions that span the V_H_ region involving well-defined regulatory segments such as the PAIR elements ^31,33,34^. To determine whether and how non-coding transcription modulates V_H_-D_H_J_H_ dynamics, we used our recently developed imaging platform to track V_H_-D_H_J_H_ motion in B cells ^27^. Specifically, we applied BCR-ABL-transformed RAG2-deficient pro-B cells, in which 240 copies of wild-type (WT) TET-operator binding sites were inserted immediately downstream of the Igh intronic enhancer and 360 copies of mutant (MUT)-TET repressor binding sites were inserted in the distal V_H_ region (Figure 1a). The WT-TET;MUT-TET;RAG2-/-pro-B cells were transduced with viruses expressing wild-type TET-repressor protein fused to GFP and mutant TET-repressor protein fused to SNAP-tag (Figure 1b). Subsequent incubation with the cell permeable dye SNAP-Cell 647-SiR and imaging with a Zeiss Elyra 7 Lattice SIM microscope platform allowed us to track V_H_-D_H_J_H_ motion in real time (Figure 1c).

**Figure 1.**
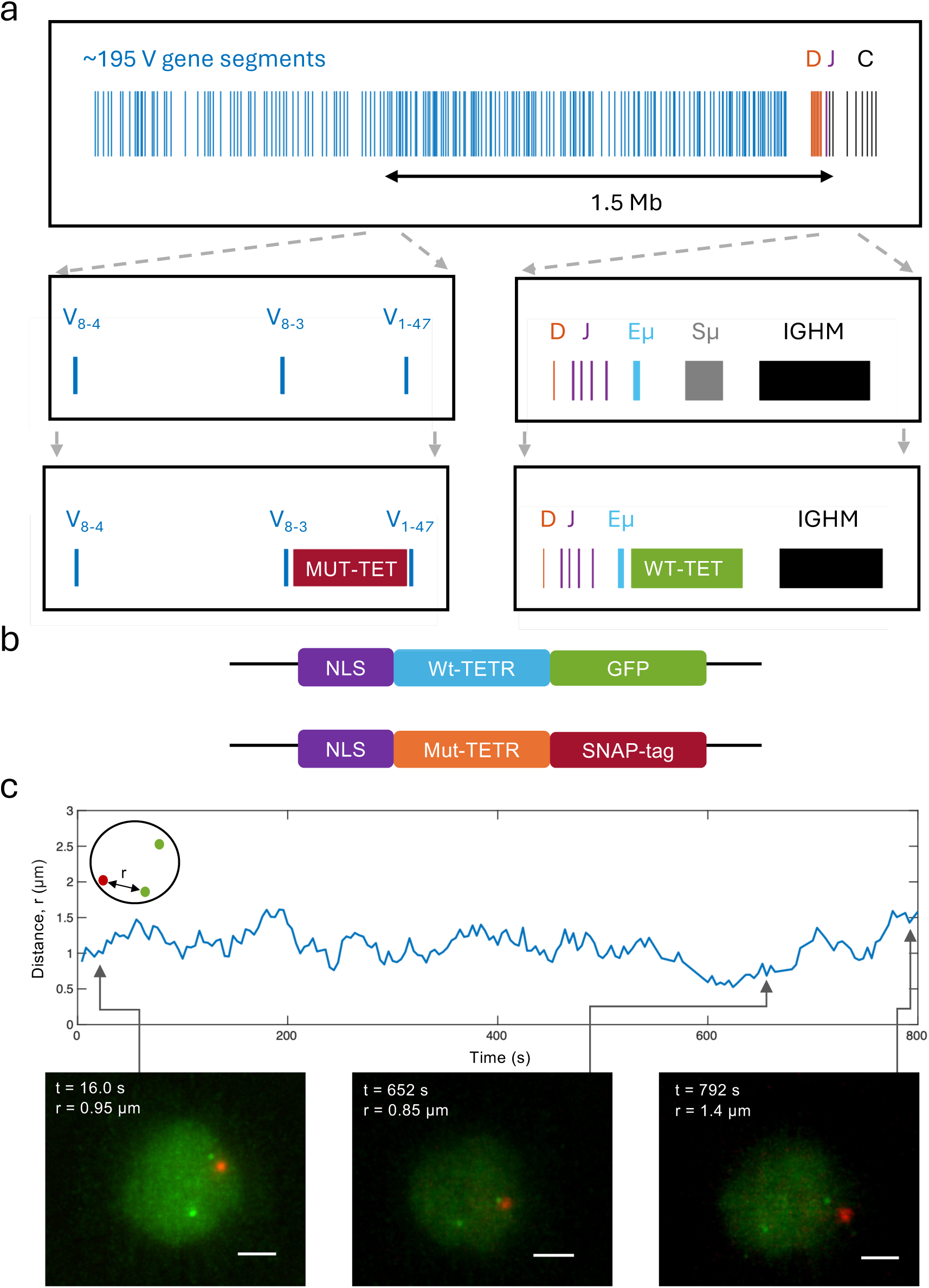
Platform to image and track V_H_-D_H_J_H_ motion in live B cells. a) Schematic of the murine Igh locus. An array of mutant-TET repressor binding sites (MUT-TETO) was inserted into the distal V_H_ region of the locus, 1.5Mb upstream of an array of WT-TET-operator binding sites (WT-TETO) inserted downstream of the D_H_J_H_ region. b) WT-TET;MUT-TET;RAG2-/-pro-B cells were transduced with viruses expressing WT-TET-GFP and MUT-TET-SNAP. c) Sample trajectory displaying the distance of separation between paired V_H_ and D_H_J_H_ regions in a single cell over time. Scale bar corresponds to 2μm.

To determine a potential role of non-coding transcription in assembling a network of crosslinks throughout the Igh locus, WT-TET;MUT-TET;RAG2-/-pro-B cells arrested in the G1 phase with STI-571 were cultured with an RNA polymerase II elongation inhibitor 5,6-dichloro-1-beta-D-ribofuranosylbenzimidazole (DRB) or, as a control, with DMSO for two hours prior to imaging (Figure 2a). Real-time PCR analysis verified that DRB suppressed both coding as well as non-coding transcription, including Iμ and PAIR4, that span the Igh locus (Figure S1b). In our previous work ^27^, such transient crosslinks were incorporated into polymer simulations to reproduce the experimentally observed chromatin dynamics and stabilization of genomic contacts. Three-dimensional (3D) V_H_ and D_H_-J_H_ genomic trajectories were recorded in DRB-treated and in DMSO-treated cells. To quantify the relative V_H_-D_H_J_H_ motion in a way that decouples it from the inevitable translational motion of the nucleus, we used the recorded 3D trajectories under each of the two conditions to calculate the mean squared displacement (MSD) as the mean squared change in the spatial distance *r* separating the intrachromosomal distal V_H_ and D_H_J_H_ regions, MSD = 〈(*r*(*t*) − *r*(*t* + *τ*))^2^〉. This approach yielded MSD of the observed apparent relative V_H_-D_H_J_H_ motion, from which we extracted the MSD of the true V_H_-D_H_J_H_ motion by correcting for nuclear rotation and for the localization error (Materials and Methods).

**Figure 2.**
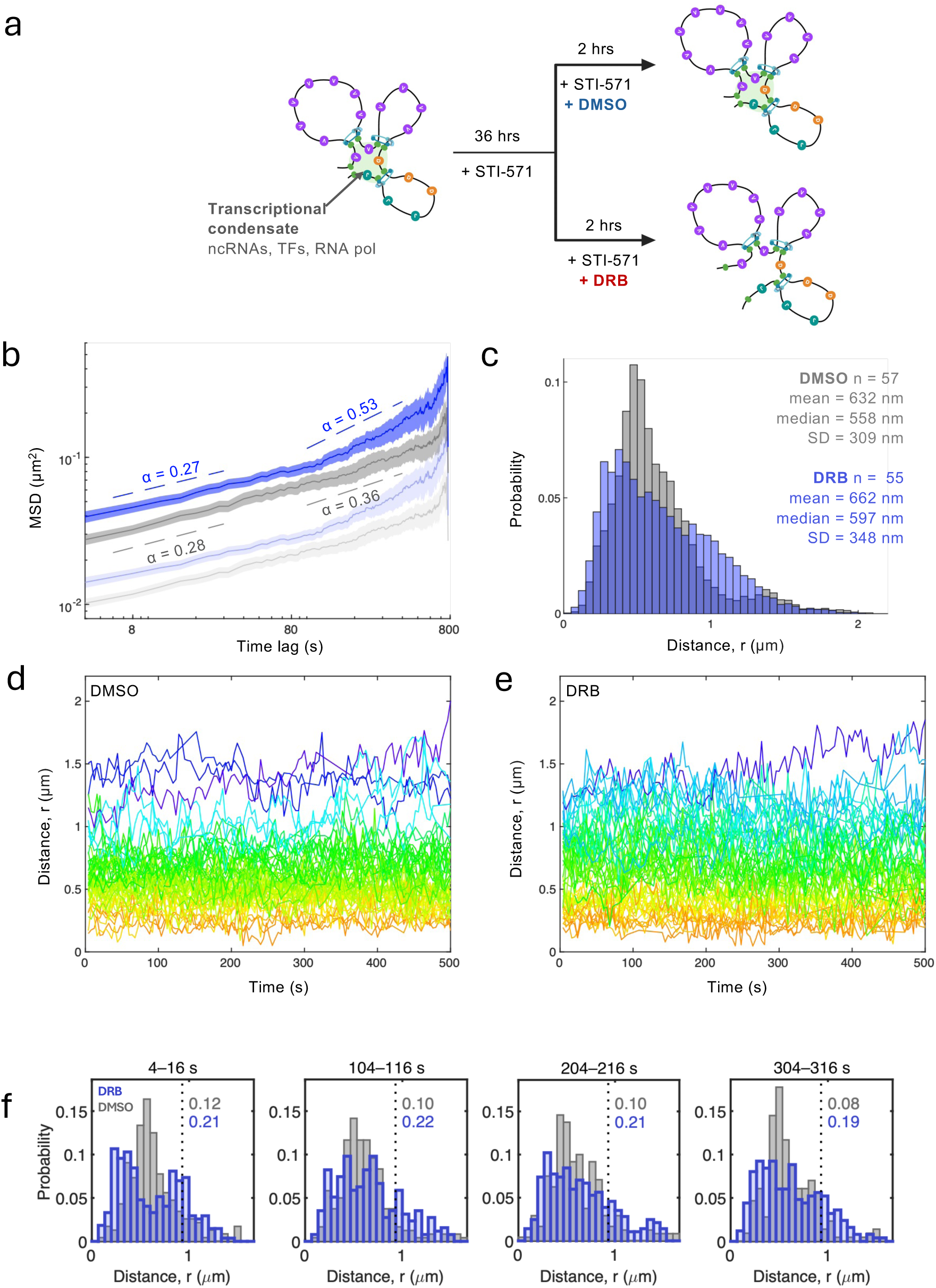
Non-coding transcription confines chromatin motion at the level of individual V_H_-D_H_J_H_ pairs. a) Experimental conditions to assess the effect of transcription on motion of remote genomic regions. Transcription inhibition was hypothesized to increase chromatin motion through loss of association of gene segments with phase separated transcriptional condensates. b) Time-and-ensemble averaged MSD of the true relative genomic motion under DRB (dark purple) and DMSO (dark grey) conditions. MSDs of observed apparent motion prior to correction for the localization error is also shown (lighter shades). c) Probability distributions of localization-error-corrected V_H_-D_H_J_H_ spatial separations over the entire imaging time under DMSO (grey) and DRB (purple) conditions. d-e) Temporal trajectories of V_H_-D_H_J_H_ distances from individual cells under DMSO (d) and DRB (e) conditions. Trajectories are color-coded based on the mean value of V_H_-D_H_J_H_ distance. The broader and less well-separated trajectories under DRB are quantified in Supplementary Fig. S7 using the normalized 90–10% range of each trajectory. f) Distributions of V_H–_D_H_J_H_ distances across time windows. Probability distributions of V_H–_D_H_J_H_ distances evaluated over short time windows (4–16 s, 104–116 s, 204–216 s, and 304–316 s) in control (DMSO, grey) and transcription-inhibited (DRB, purple) cells. Distributions in DMSO are similar across time windows and remain relatively compact. In contrast, DRB exhibits broader distributions with enhanced tails, indicating increased heterogeneity and more frequent large excursions. The numbers in each panel indicate the fraction of distances exceeding a fixed threshold (defined as the 90th percentile of the pooled DMSO distribution across all windows), quantifying the relative weight of the distribution tails. A per-cell analysis using the same time windows shows an approximately twofold increase in the fraction of distances exceeding the threshold in DRB compared to DMSO (mean 0.197 vs 0.103; permutation test p ≈ 0.051), consistent with the pooled distributions. These results are consistent with a reduction of local confinement upon transcription inhibition.

The time-and-ensemble averaged MSD revealed that inhibition of transcription elongation significantly increased average V_H_-D_H_J_H_ motion over relatively long times (> 10 minutes) (Figure 2b). While V_H_-D_H_J_H_ motion in control cells (DMSO) was found to be strongly subdiffusive (scaling exponent α≈0.3), consistent with our previous work ^27^, interference with transcription elongation increased the population-averaged scaling exponent from α = 0.36 to α = 0.53 for intermediate and long-time scales, with a modest effect on the diffusion coefficient (Table S1a). Furthermore, inhibition of transcription elongation resulted in an increase for both the median and variance of the distribution of V_H_-D_H_J_H_ spatial separations (Figure 2c).

While the MSD revealed a significant increase in the average V_H_-D_H_J_H_ motion following inhibition of transcription elongation, by virtue of being an ensemble-averaged quantity it masked the effect of transcription on the motion of individual V_H_-D_H_J_H_ pairs. In order to gain insights into chromatin dynamics beyond the population average, we examined the temporal V_H_-D_H_J_H_ distance trajectories in individual cells. Consistent with our previous findings, the trajectories in the control cells (DMSO) fluctuated around their respective means that remained nearly constant over time. This behavior manifested itself as the distinct rainbow-like pattern where the trajectories color-coded based on the mean value appeared “demixed” from one another (Figure 2d) ^27^. Notably, however, V_H_-D_H_J_H_ trajectories in cells that were cultured under conditions that inhibit transcription elongation (DRB) showed loss of the de-mixing effect (Figure 2e). The key insight from the observed change in the dynamics is that transcription constrains V_H_-D_H_J_H_ spatial distances and the extent of fluctuations not just as an average effect across the population of cells but also within each V_H_-D_H_J_H_ pair.

To quantify the reduced demixing visible in the trajectory plots (Figure 2d–e), we calculated the width of distance fluctuations for each V_H–_D_H_J_H_ pair relative to its own typical separation (Figure S2a). Specifically, we computed the normalized 90–10% range, (q90 − q10) / median(r), where r(t) is the distance trajectory and q10 and q90 are its 10th and 90th percentiles over the time interval shown. This metric was significantly increased under transcription inhibition (DRB) compared to control (DMSO) (0.537 vs 0.439; permutation test p = 0.0046), indicating that individual V_H–_D_H_J_H_ pairs explore a broader range of separations. An independent measure, the coefficient of variation std(r) / mean(r), yielded consistent results (0.212 vs 0.176; p = 0.006) with large early-to-late excursions (Figure S2b). These analyses provide a quantitative measure of the reduced demixing and support a model in which nascent transcription establishes local confinement that stabilizes interactions between spatially proximal genomic elements.

We previously showed that the de-mixing effect in the distance trajectories is consistent with local spatial confinement arising from transient crosslinks within the chromatin fiber. To test whether nascent transcription is a primary mechanism underlying this local confinement, we examined the origin of the increased spread in the distance distribution upon transcription inhibition (Figure 2c). Specifically, we analyzed the distributions of V_H–_D_H_J_H_ distances over short time windows (Figure 2f). In control cells (DMSO), the distributions are similar across time windows and remain relatively compact, consistent with stable local confinement. In contrast, in DRB-treated cells, the distributions become broader and exhibit enhanced tails, indicating increased heterogeneity and more frequent large excursions in spatial separation away from locally proximal configurations. In the framework introduced in our previous work ^27^, transient chromatin crosslinks act as local constraints that stabilize genomic contacts and suppress large fluctuations in spatial separation. Thus, the broadened distributions and enhanced tails observed upon transcription inhibition are consistent with weakening of this local confinement. This behavior indicates that transcription establishes local confinement, limiting chromatin segments to a narrower range of spatial separations. Quantification of the distribution tails showed an approximately twofold increase in the fraction of large-distance events in DRB compared to DMSO (Figure 2f). A per-cell analysis yielded a similar increase (mean 0.197 vs 0.103; p = 0.05). These results are consistent with the observed broadening of the distributions and increased tail weight under transcription inhibition. Taken together, these results support a model in which nascent transcription acts to establish local confinement that stabilizes interactions between spatially proximal genomic elements.

### Architectural protein CTCF constrains population-average chromatin dynamics

To determine the effect of CTCF on chromatin dynamics, we studied V_H_-D_H_J_H_ motion in cells that were depleted of CTCF expression. To accomplish this objective, we bi-allelically inserted the FKBP12^F36V^ degron and the fluorescent marker mScarlet into the CTCF locus using the strategy described previously for neutrophil progenitors (Figure 3a). Correct targeting was verified by genotyping. WT-TET;MUT-TET;RAG2-/-pro-B cells carrying CTCF degrons on both alleles were then transduced with viruses expressing WT-TET-GFP and MUT-TET-SNAP-tag. Transduced cells were then incubated with the cell-permeable ligand dTAG-13 to induce CTCF degradation (Figure 3a). Flow cytometric analysis confirmed that 6 hours of dTAG-13 treatment substantially reduced CTCF abundance (Figure 3b). We next examined how CTCF depletion affects V_H_-D_H_J_H_ chromatin dynamics. We found that CTCF depletion resulted in a modest increase in V_H_-D_H_J_H_ motion, as manifested by a small but statistically significant increase in both the anomalous scaling exponent and diffusion coefficient extracted from the MSD (Figure 3c and Table S1a). Consistent with this result, the probability distributions of the spatial distances between V_H_ and D_H_J_H_ elements were similar for the two conditions, with only a modest increase in the median spatial distance (Figure 3d). Examining V_H_-D_H_J_H_ distance trajectories in single cells further revealed that CTCF depletion did not interfere with the local character of chromatin motion, preserving the rainbow-like pattern in color-coded trajectories (Figures 3e-f). A single DMSO trajectory exhibited a much larger spatial separation (> 3 μm, Figure 3d) than the remaining trajectories (Figure S3), illustrating the inherent heterogeneity and stochastic nature of V_H_-D_H_J_H_ motion even under basal conditions. Together, these results indicate that CTCF occupancy modestly constrains the population-average statistical properties of V_H_-D_H_J_H_ dynamics but has little effect on their spatiotemporal organization.

**Figure 3.**
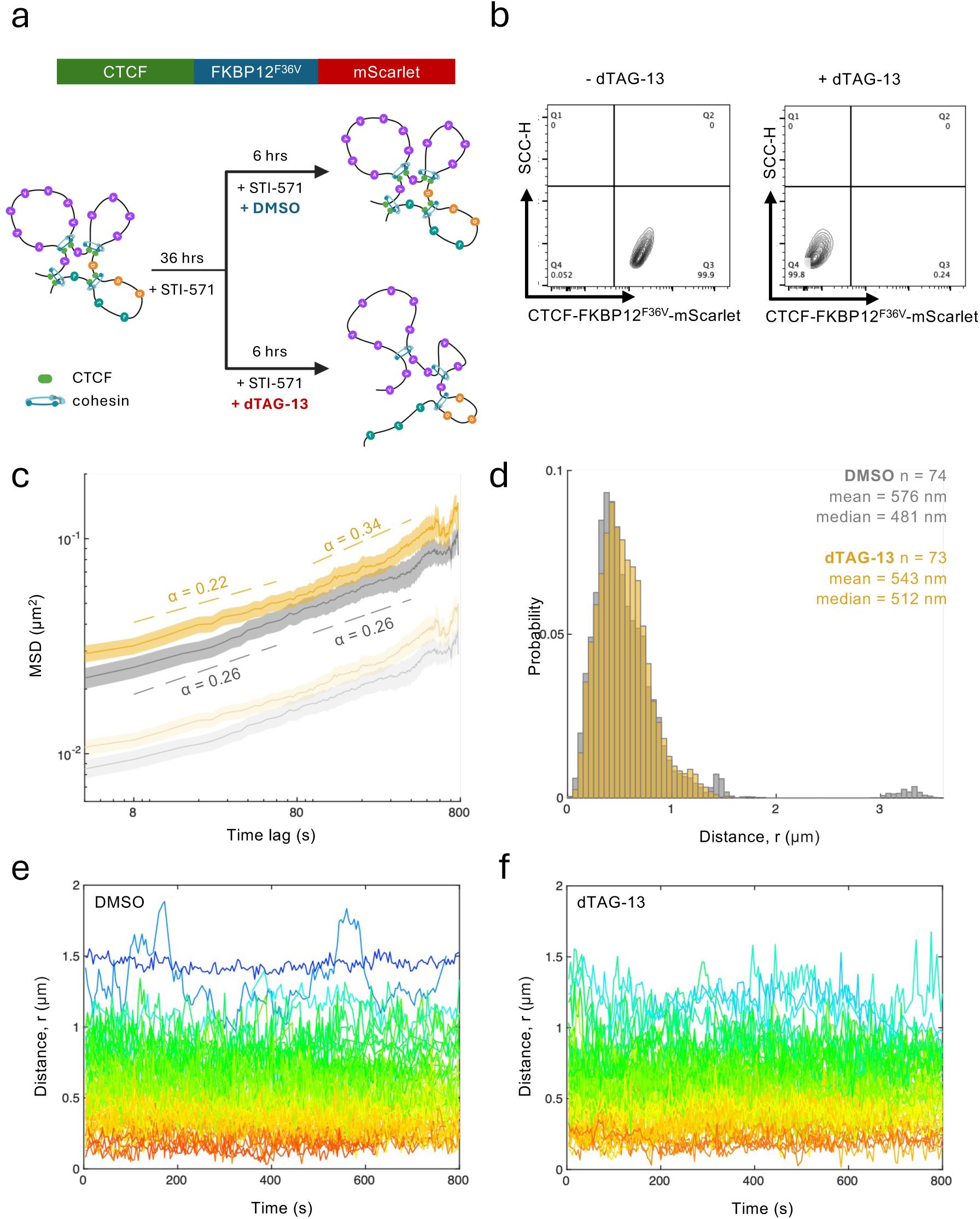
Architectural protein CTCF constrains population-average chromatin motion. a) Experimental conditions to assess the effect of CTCF on motion of remote genomic regions. CTCF was depleted by culturing WT-TET;MUT-TET;RAG2-/-;CTCF-mScarlet-FKBP12^F36V^-mScarlet pro-B cells with dTAG-13. CTCF depletion is known to cause the assembly of loops to proceed past their normal boundaries. Top panel indicates CTCF boundary elements (green oval dots) in wild-type pro-B cells. The lower panel shows the absence of boundary elements in CTCF-depleted pro-B cells, allowing loop extrusion to proceed beyond the normal boundaries. b) mScarlet expression following 6 hrs DMSO and dTAG-13 treatment of CTCF-FKBP12^F36V^-mScarlet cells. c) Time-and-ensemble averaged MSD curves of relative genomic motion under DMSO (dark grey) and dTAG-13 (dark yellow) conditions. MSD curves prior to error correction are also shown (light shades). d) Probability distributions of V_H_-D_H_J_H_ spatial separations over the entire imaging time under dTAG-13 and DMSO conditions. e-f) Temporal trajectories of V_H_-D_H_J_H_ distances from individual cells under DMSO (e) and dTAG-13 (f) conditions. Trajectories are color-coded based on the mean value of V_H_-D_H_J_H_ distance.

### Cohesin-mediated loop extrusion provides large-scale confinement that co-localizes V_H_ and D_H_J_H_ elements and constrains average dynamics

To determine the effect of cohesin-mediated loops on V_H_-D_H_J_H_ motion, we bi-allelically tagged RAD21 with FKBP12^F36V^ and mScarlet in BCR-ABL transformed WT-TET;MUT-TET;RAG2-/-pro-B cells (Figure 4a). Bi-allelic insertion was confirmed by genotyping (data not shown). Subsequently, WT-TET;MUT-TET;RAG2-/-pro-B cells carrying RAD21-degrons were transduced with viruses expressing WT-TET-GFP and MUT-TET-SNAP-tag, and cultured in the presence of STI-571 and dTAG-13 to induce RAD21 degradation (Figure 4b). Western blotting confirmed efficient RAD21 degradation following dTAG-13 treatment (Figure S4). Temporal trajectories of V_H_-D_H_J_H_ spatial distances were measured in the cells cultured with and without dTAG-13 and time-and ensemble-averaged MSDs for the two conditions were calculated (Figure 4c). MSD analysis revealed a substantial increase in both the diffusion coefficient and the scaling exponent in pro-B cells depleted of RAD21 (Table S1a). RAD21 depletion also resulted in an increase in the average V_H_-D_H_J_H_ spatial distance (Figure 4d).

**Figure 4.**
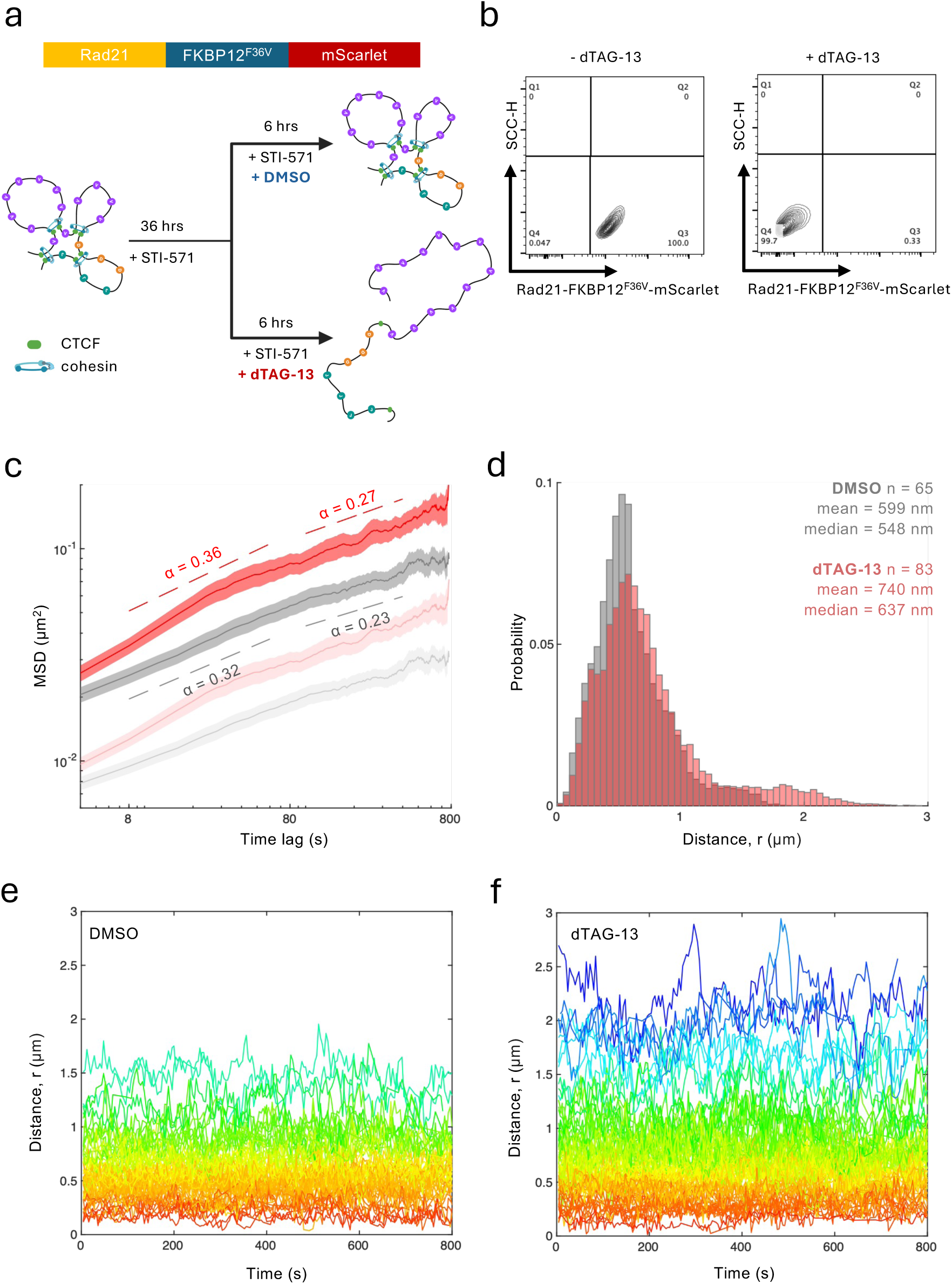
The assembly of cohesin-mediated loops bring paired V_H_ and D_H_J_H_ segments into spatial proximity and restrain V_H_-D_H_J_H_ motion. a) Experimental conditions to assess the effect of cohesin-mediated loops on motion of remote genomic regions. WT-TET;MUT-TET;RAG2-/-;RAD21-FKBP12^F36V^-mscarlet pro-B cells were treated with dTAG-13 to deplete endogenous RAD21 expression and disrupt the assembly of cohesin-mediated chromatin loops. b) mScarlet expression is abrogated following 6 hrs dTAG-13 treatment in RAD21-FKBP12^F36V^ -mScarlet cells. c) Time-and-ensemble averaged MSD curves of true relative genomic motion from RAD21-FKBP12^F36V^-mScarlet cells treated with DMSO (dark grey) and dTAG-13 (dark red). MSD curves prior to error correction are also shown (light shades). d) Probability distributions of V_H_-D_H_J_H_ distances under DMSO (grey) and dTAG-13 (red) conditions. e-f) Temporal trajectories of V_H_-D_H_J_H_ distances from individual cells under DMSO (e) and dTAG-13 (f) conditions. Trajectories are color-coded based on the mean value of V_H_-D_H_J_H_ distance.

To unravel the mechanistic origin of the observed increase in MSD upon removal of RAD21, we examined the individual V_H_-D_H_J_H_ trajectories under the two conditions (Figures 4e and 4f). The trajectories revealed the effect of RAD21 depletion that was masked by the ensemble-averaging in the MSD: the emergence of a sub-population of V_H_-D_H_J_H_ pairs with relatively large spatial distances as well as large distance fluctuations (Figure 4f). In the rest of the population, the de-mixing effect was preserved, as manifested by the rainbow-like pattern adopted by color-coded trajectories (Figure 4f). These two sub-populations explain the bimodal distribution of V_H_-D_H_J_H_ spatial distances observed following RAD21 depletion (Figure 4d). These analyses show that the increase in average V_H_-D_H_J_H_ motion following RAD21 depletion arises from the emergence of a highly mobile sub-population of V_H_-D_H_J_H_ trajectories. Together, these results indicate that cohesin-mediated chromatin loops provide a global-scale confinement that brings V_H_ and D_H_J_H_ elements into spatial proximity and constrains average V_H_-D_H_J_H_ motion.

### Cohesin-mediated loop extrusion reinforces self-similarity of V_H_-D_H_J_H_ motion across temporal/spatial scales

The results of the RAD21 depletion described above revealed a newly-emerged sub-population of chromatin configurations that were highly mobile as well as characterized by relatively large spatial distances separating paired V_H_ and D_H_J_H_ elements (Figure 4c-f). To quantitatively define this sub-population, a threshold of 1.25 μm corresponding to the position of the valley in the distance distribution was used (Figure 4d). The sub-population with r > 1.25 μm accounted for 4.6% of cells expressing wild-type levels of RAD21 and increased to 12.0% following RAD21 depletion (Figure 4d). The trajectories in this newly emerged sub-population exhibited substantially larger excursions around their mean values than the excursions observed in the cells that expressed wild-type levels of RAD21 (Figure 5a). The increase in mobility was also evident in the individual time-averaged MSDs as well as the time-and-ensemble averaged MSD for the emerged sub-population of trajectories (Figure 5b). This sub-population was further found to account for most of the difference in the population-wide MSD between the cells depleted for RAD21 and cells that expressed wild-type levels of RAD21 (Figure 5c). Furthermore, the MSD for the emerged sub-population exhibited a rather steep slope (scaling exponent α = 0.54) over short time lags followed by a saturation (α = 0.14) at a value of 0.28 μm^2^ over intermediate-to-long time lags (Figure 5c; Table S1). These findings indicate that paired V_H_-D_H_J_H_ elements in the newly emerged sub-population behave as a free polymer on short time scales (the scaling exponent is close to the noninteracting Rouse chain value of α = 0.5), eventually reaching a large-scale spatial confinement (the scaling exponent approaches zero) (Figure 5c). Collectively, these observations suggest that, in cells with diminished loop extrusion activity, a subset of paired V_H_-D_H_J_H_ elements no longer experienced local confinement but remained constrained on a larger scale.

**Figure 5.**
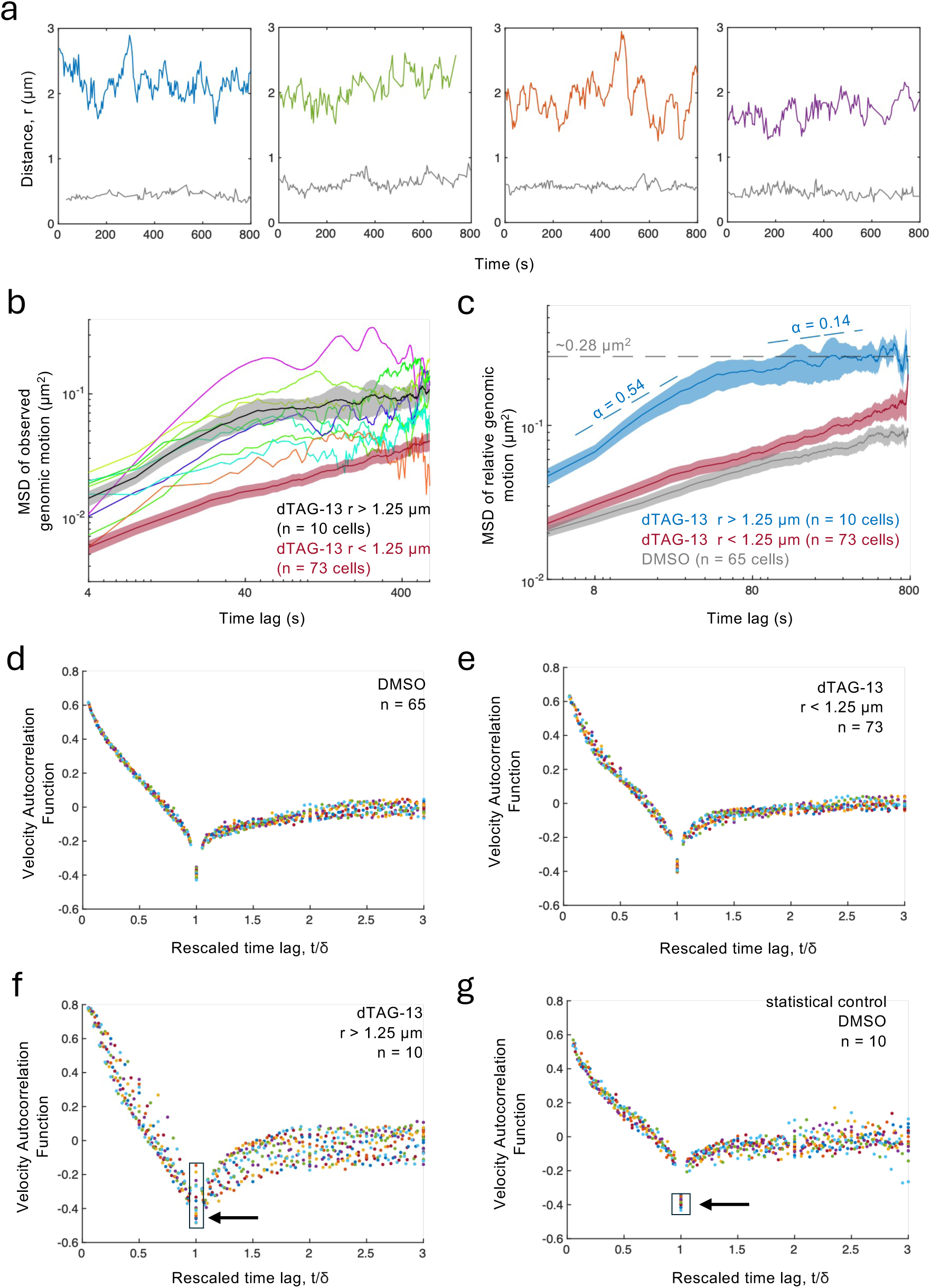
Cohesin-mediated loops preserve self-similarity of V_H_-D_H_J_H_ motion across temporal/spatial scales. a) Representative trajectories from dTAG-13-treated RAD21-FKBP12^F36V^-mScarlet cells with large spatial V_H_-D_H_J_H_ separations (r > 1.25 μm). Trajectories from DMSO treated cells are shown in grey. b) Thin colored lines are individual time-averaged MSD curves for the sub-population of trajectories with large spatial separations (r > 1.25 μm) from RAD21-depleted cells. Also shown are the corresponding time-and-ensemble averaged MSDs for the trajectories with large spatial separations (r > 1.25 μm) (grey) and with normal separations (r < 1.25 μm) (red). c) Time-and-ensemble averaged MSD curves of relative genomic motion for dTAG-13 trajectories with high spatial separations (r > 1.25 μm) (blue), dTAG-13 trajectories with normal separations (r < 1.25 μm) (red), and DMSO trajectories (grey). d) Velocity autocorrelation functions (VAFs) computed from trajectories in DMSO-treated Rad21-FKBP12^F36V^-mScarlet cells collapse onto a single curve upon rescaling of the time axis, indicating self-similarity. e) Self-similarity revealed by the rescaled VAFs computed from the trajectories with normal spatial separations (r < 1.25 μm) in dTAG-13-treated Rad21-FKBP12^F36V^-mScarlet cells. f) Loss of self-similarity revealed by the rescaled VAFs computed from n=10 trajectories with large spatial separations (r > 1.25 μm) in dTAG-13-treated Rad21-FKBP12^F36V^-mScarlet cells. g) A statistical control shows rescaled VAFs for 10 randomly sampled trajectories from DMSO-treated Rad21-FKBP12^F36V^-mScarlet cells.

To determine whether this newly emerged dynamical state retained the scale-free behavior observed in wild-type cells, we analyzed the velocity autocorrelation functions computed from the genomic trajectories that were recorded in the presence and absence of RAD21. To obtain the velocity autocorrelation functions, we calculated the relative velocity of a given genomic pair as a change in V_H_-D_H_J_H_ distance during the time interval δ divided by that time interval, and then computed the correlation between two such velocities temporally separated by the interval τ. To probe the motion across a range of time scales, the correlations were examined for different values of δ and (for each δ) over a range of values of τ. We previously showed that the velocity autocorrelation functions for intrachromosomal distal V_H_ and D_H_J_H_ regions exhibited negative values that approached the theoretical “extreme confinement” limit of -0.5 ^27^. Negative correlations indicate that the genomic regions experience a pushback from their environment – a restoring force whereby motion in one direction is likely to be followed by motion in the opposite direction. Such anti-persistent behavior was shown to be a consequence of diffusion in a viscoelastic environment ^27^.

The velocity autocorrelation functions for the cells with wild-type levels of RAD21 revealed negative correlations, as well as self-similarity manifested as the collapse onto a single curve upon rescaling of the time axis (Figure 5d), indicating similar patterns of motion across temporal and spatial scales, consistent with previous observations ^27^. Likewise, self-similarity was observed for the V_H_-D_H_J_H_ trajectories with spatial separations r <1.25 μm in the cells depleted of RAD21 (Figures 5e). Notably, however, the velocity autocorrelation functions for the trajectories with large spatial separations, r >1.25 μm, in cells depleted of RAD21, displayed a significant scatter upon rescaling of the time axis, indicating a loss of self-similarity (Figure 5f). To rule out the possibility that the scatter in the rescaled data was due to the small sample size (n=10), we performed multiple statistical controls by calculating the velocity autocorrelation functions using n=10 trajectories randomly sampled from cells with wild-type levels of RAD21 (DMSO) (Figure 5g and Figure S5). This approach confirmed that the loss of self-similarity observed for V_H_-D_H_J_H_ trajectories with large spatial separations is an intrinsic property of this sub-population. Thus, the loss of self-similarity of V_H_-D_H_J_H_ motion is associated with the crossover between short-time free-polymer-like dynamics and long-time confinement-dominated dynamics as revealed by MSD. The coexistence of these distinct dynamical regimes introduces characteristic temporal and spatial scales into the trajectories, leading to the breakdown of the rescaling collapse observed for self-similar motion. Together, these results indicate that, in a subset of cells (12.0%), paired V_H_ and D_H_J_H_ gene segments whose spatial proximity depends on cohesin-mediated loop extrusion enter a distinct dynamical state characterized by (i) enhanced mobility, (ii) loss of self-similarity, that is, loss of similar patterns of motion across temporal and spatial scales, and (iii) a crossover from free-polymer dynamics to dynamics dominated by a large-scale spatial confinement.

### The frequency of V_H_-D_H_J_H_ encounters is governed by the interplay of chromatin diffusivity and spatial confinement

The effect of each of the perturbations described above on the frequency, *f*, of V_H_-D_H_J_H_ encounters can be assessed by estimating the fold change in the mean time for the first encounter, or mean first passage time (MFPT=1/*f*), using dimensional analysis ^35^. From the dimensions of MSD, [MSD] = [length]^2^ = [*D*] [time]^α^, we have [time] = [length]^2/α^ [*D*]^-1/α^, where *D* is the diffusion coefficient and α is the anomalous scaling exponent. The relevant parameter with the dimensions of length in this first-passage-time problem is the radius of confinement, *R*. The size of the spatial confinement can be estimated from the measured median V_H_-D_H_J_H_ separation, while the values of *D* and α can be extracted directly from the time-and-ensemble-averaged MSD (Table S1). For any two experimental conditions, 1 (control) and 2 (perturbation), the above dimensional analysis argument yields the fold-change in MFPT upon the perturbation of interest (Figure 6a):

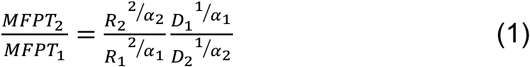

**Figure 6.**
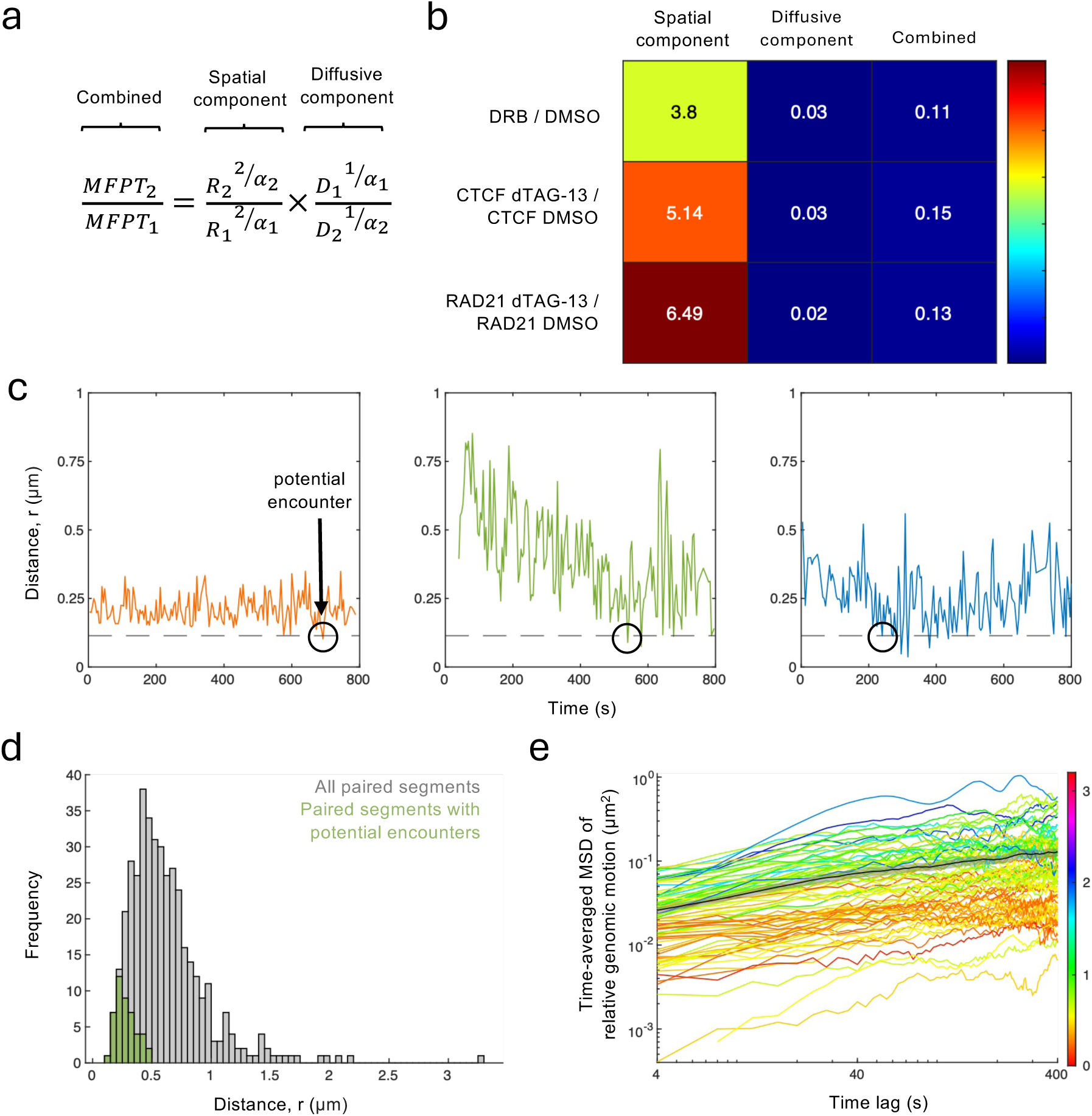
The frequency of V_H_-D_H_J_H_ encounters is governed by the interplay of chromatin diffusivity and spatial confinement. a) Equation obtained by dimensional analysis to estimate the fold-change in MFPT upon perturbation of interest. b) Fold changes in MFPT for each perturbation, along with individual spatial and diffusive components. c) Sample trajectories with potential encounters (circles) between V_H_ and D_H_J_H_ genomic regions. d) Probability distributions of V_H_-D_H_J_H_ spatial distances for all paired regions (grey) and for the subset with a potential encounter (green), showing that potential encounters occur almost exclusively among spatially proximal (separated by less than 500 nm) paired V_H_-D_H_J_H_ elements. e) MSD curves averaged over time (but not over ensemble) computed from individual trajectories in RAD21-mScarlet-FKBP12^F36V^ cells treated with dTAG-13. The MSD curves are color-coded according to the heatmap on the right based on the mean value of the V_H_-D_H_J_H_ distance in the corresponding trajectory, revealing an overall correlation between chromatin mobility and spatial separation.

Equation (1) reveals that the fold-change in MFPT, or equivalently the inverse of the fold-change in the encounter frequency, can be viewed as a superposition of the spatial component (governed by the change in the confinement size *R*) and the diffusive component (governed by the change in the diffusion coefficient *D*), with both components additionally regulated by the change in the scaling exponent α (Figure 6a). The spatial and diffusive components were obtained by separately evaluating the confinement-dependent (R) and diffusivity-dependent (D) terms in Eq. (1), with both contributions modulated by the anomalous scaling exponent α (Figure 6b).

With the values of *D*, *α* and *R* extracted from the experimental data for each of the perturbation experiments, Equation (1) revealed a general trend (Figure 6b). All the introduced perturbations loosened V_H_-D_H_J_H_ confinement (spatial component >1) and increased V_H_-D_H_J_H_ motion (diffusive component < 1; note the inverse of the ratio of the diffusion coefficients in Eq. (1)). Further, all the perturbations were predicted to reduce the mean waiting times for the first encounter (the combined effect on the two components in the MFPT fold-change <1) and thus increase the encounter frequency (Figure 6b).

Interestingly, the fold-change in V_H_-D_H_J_H_ MFPT exhibited the same trend as that of the diffusive component (Figure 6b), suggesting that, among genomic regions already positioned within close spatial proximity, V_H_-D_H_J_H_ diffusivity, reflecting the ability of chromatin segments to explore space through motion, amplified by the inverse of the anomalous scaling exponent (D^1/α^), primarily modulates the frequency of interactions between paired regulatory and coding elements. Indeed, even a modest increase in V_H_-D_H_J_H_ motion, as revealed by CTCF-depleted trajectories (Table S1), substantially reduced the waiting time for V_H_-D_H_J_H_ encounters (Figure 6b).

To gain intuition about the trend revealed by the dimensional analysis argument, we focused on the trajectories of spatial distances in which potential encounters between V_H_ and D_H_J_H_ elements occurred within the imaging time (Figure 6c). All the V_H_-D_H_J_H_ trajectories marked by potential genomic encounters had a mean spatial distance below 500 nm (Figure 6d), indicating a critical role for spatial proximity.

Notably, we found that the paired genomic regions that were positioned in close spatial proximity (r <500 nm) were significantly less mobile than those with larger spatial separations (r >500 nm), across all experimental conditions (Figures S6). As further evidence of this correlation, individual time-averaged MSDs for RAD21-depleted trajectories, color-coded according to the corresponding mean spatial distance, exhibited substantial segregation based on spatial proximity (Figure 6e). The observed segregation demonstrates – on the level of individual V_H_-D_H_J_H_ pairs – that chromatin motion and the extent of spatial proximity are closely linked. Taken together, the dimensional analysis of MFPTs combined with experimentally determined parameter values and direct examination of the trajectories indicates that (i) potential encounters require spatial proximity (r < 500 nm) of the genomic elements in the folded chromosome, and (ii) for the elements that satisfy the proximity requirement, the frequency of encounters is primarily regulated by chromatin diffusivity with further amplification by the subdiffusive scaling exponent.

## Discussion

It is now established that the motion of paired genomic regions in eukaryotes is strongly subdiffusive. We have previously proposed that the subdiffusive motion is imposed by a network of reversible crosslinks plausibly involving epigenetic marks, chromatin remodelers, transcription factors, and chromatin architectural proteins. Here we show that, while both transcription and cohesin-mediated chromatin loops confine distally located V_H_ and D_H_J_H_ gene segments in the Igh locus, they do so in different ways. Transcription imposes a local-scale confinement on the distal V_H_ and D_H_J_H_ regions that have been brought into spatial proximity. We found that, consistent with previous studies, transcription inhibition increased chromatin motion without significantly increasing the spatial separation of paired V_H_-D_H_J_H_ elements ^36^. Theory and simulations have previously suggested that a weakly-crosslinked polymer (weak gel) model not only reproduced the average behavior of V_H_-D_H_J_H_ pairs but also the pattern of motion of individual gene pairs ^27^. Here, we found that the effects of transcription on chromatin dynamics reproduced the effects of crosslinks. We propose that nascent transcription constrains the relative motion of paired V_H_-D_H_J_H_ gene segments through engagement with phase-separated transcription factories, each involving a dense network of chromatin, transcription factors, and transcriptional co-activators ^15,37^. Motion of genomic regions within this dense network is naturally constrained, whereas disengagement of the segments from the transcriptional condensates in the absence of transcription results in increased mobility.

While it has previously been reported that loss of cohesin increases chromatin motion, our findings indicate that cohesin restricts chromatin motion across much larger genomic distances (∼1.5 Mb) than previously shown (∼150 kb) ^19,38^. We also found that RAD21-depletion enriched for a subset of trajectories (∼12%) that spanned large spatial distances. This raises the question of how distinct pools of V_H_-D_H_J_H_ trajectories in pro-B cells that are depleted for loop extrusion arise. Previous studies have shown that pro-B cells depleted for loop extrusion are associated with a substantial decline in local genomic interactions and adopt a plaid-like pattern in chromosome conformation capture maps ^12^. It has been proposed that such plaid-like patterns arise because, in the absence of loop extrusion, TAD density across the chromatin fiber declines, leading to enhanced chromatin mobility ^6^. Increased mobility, in turn, allows for remote genomic regions sharing similar epigenetic identities. Enrichment for such long-range interactions ultimately yields more pronounced and segregated euchromatic and heterochromatic compartments. Hence, we propose that loop extrusion in pro-B cells suppresses the emergence of highly mobile V_H_-D_H_J_H_ trajectories and the associated loss of spatial confinement, thereby limiting the assembly of highly segregated euchromatic and heterochromatic compartments. Taken together, our findings support a hierarchical picture of “confinement within confinement,” in which transcription-dependent local confinement is nested within the larger-scale confinement established by cohesin-mediated loops.

Immunoglobulin heavy chain locus rearrangement is regulated by yet another loop extrusion factor, WAPL. WAPL expression is down-regulated in early B cell progenitors to enrich for distally-located V_H_-D_H_J_H_ rearrangements ^38,39^. It will be interesting to determine whether and how WAPL regulates V_H_-D_H_J_H_ mobility and Igh locus confinement to instruct V_H_-D_H_J_H_ rearrangement.

What is the functional advantage to having a hierarchy of confinements? We suggest that the global confinement prepares the ground for genomic encounters by exploiting a general physical property of subdiffusive motion – strong enhancement of search efficiency upon confining the space. On a much smaller scale, the local confinement tunes the properties of the immediate neighborhood of gene segments in a tightly regulated manner to facilitate interactions between the segments trapped within and insulate those left outside. Thus, the global confinement of cohesin-mediated chromatin loops brings a subset of V_H_-gene segments from across the locus into spatial proximity, poising them for potential encounters with D_H_J_H_ elements. In turn, the local confinement mediated by transient crosslinks, whose assembly is tightly regulated during developmental progression, stabilizes interactions within this subset of spatially proximal segments while insulating them from the rest of the locus, and subsequently disassembles to allow large-scale reconfiguration of loops that span the Igh locus such that a new subset of V_H_-segments is poised for potential encounters. In this manner, the hierarchical nature of spatial confinement provides the balance of stability and mobility that is necessary to diversify genomic encounters, orchestrate their timing, and regulate their frequencies to ultimately generate a diverse antibody repertoire.

We note several limitations of the present study. First, our measurements rely on tandem arrays of Tet operator binding sites and TET repressors. Previous work has demonstrated that insertion of TET operator arrays into the Igh locus does not interfere with B-cell maturation or Igh locus rearrangement ^26^. Moreover, TET repressors lack domains known to promote extensive oligomerization, long-range genomic interactions, or the formation of nuclear condensates, making it unlikely that they directly perturb loop extrusion or chromatin confinement. Nevertheless, such effects cannot be formally excluded. Second, transcription was inhibited using the RNA polymerase II elongation inhibitor DRB. Although the treatment was brief, indirect effects resulting from altered expression of factors involved in chromatin organization cannot be excluded. Future studies involving systematic perturbation of non-coding transcripts across the Igh locus should help identify the transcripts responsible for regulating local chromatin motion. Finally, although flow cytometry confirmed efficient depletion of CTCF and RAD21 by culturing the cells in the presence of dTAG-13, we cannot exclude the possibility that residual chromatin-bound CTCF or RAD21 persisted in a subset of cells.

One of the central observations of this study was the strong link between the extent of V_H_-D_H_J_H_ motion and spatial proximity. Indeed, the same trend persisted both at the level of individual V_H-DHJH t_rajectories and at the population level, and under all experimental conditions: more spatially proximal genomic regions exhibited less motion. At the same time, we found that, while paired V_H_-D_H_J_H_ interactions require spatial proximity, the frequencies of such interactions were dominated by chromatin diffusivity. The established link between spatial separation of V_H_ and D_H_J_H_ elements and V_H_-D_H_J_H_ motion suggests that spatial proximity may serve not only to facilitate genomic interactions but also to restrict them. We therefore suggest that the generation of a diverse antibody repertoire depends on striking a balance between the freedom of V_H_-D_H_J_H_ motion and spatial proximity of V_H_ and D_H_J_H_ elements, a balance achieved through the hierarchy of confinements imposed by transcription and cohesin-mediated loop extrusion.

## Online Methods

### Cell culture

Murine BCR-ABL-transformed RAG2-deficient pro-B cells carrying WT-TET and MUT-TET operator sites have been described in earlier studies ^27^. Cells were cultured in either Optimem or phenol free Optimem with 10% fetal calf serum, antibiotics, glutamine and β-mercaptoethanol. Cell culture media was supplemented with 2 ng/mL stem cell factor and 10 ng/mL interleukin-7. STI-571 treatments were performed at a concentration of 10 μM for a minimum of 36 hours prior to imaging. dTAG-13, was supplemented to media at a concentration of 0.5 μM.

### Viral Production and Transduction

TetR-eGFP and mut-TetR-SNAP constructs were previously published ^26,27^. Vectors were grown up and isolated with either CsCl purification or with Genejet Endo-free Plasmid Maxiprep Kit. Viral vectors were transfected into 293T cells by calcium phosphate transfection, along with a retroviral packing plasmid as described ^42^. Media was changed to fresh media 1 day post transfection, and virus containing media was harvested 2 and 3 days post transfection. Virus containing media was stored at -80C. Pro-B cells were transduced via spin transfection for 1 hr at 2000 rpm and 37C.

### Interfering with transcriptional elongation

To block transcriptional elongation cells were treated with 100 μM 5,6-Dichloro-1-β-D-ribofuranosylbenzimidazole (DRB) for 2 hours prior to imaging. RNA was isolated and quantified using a NanoDrop spectrophotometer. One microgram of total RNA was reverse transcribed into cDNA using a Bio-Rad iScript cDNA Synthesis Kit (Bio-Rad 1708891) according to the manufacturer’s protocol. Quantitative PCR was performed using SsoAdvanced Universal SYBR Green Supermix (Bio-Rad 1725270) on a CFX96 Touch Real-Time PCR Detection System (Bio-Rad). Relative gene expression was calculated using the comparative Ct (2^-DDCt) method. Ct values were normalized to the DMSO/STI-571 treated control samples, and expression levels were calculated relative to the indicated control sample. All reactions were performed in technical triplicates. Data are presented as mean ± standard deviation (SD) from at least two independent biological replicates. Statistical analyses were performed using GraphPad Prism software.

### Western blotting

BCR-ABL transformed RAD21-FKBP12F36V-mscarlet pro-B cells carrying WT-TET and MUT-TET operator sites were cultured in the presence or absence of dTAG-13 for 6 hours. Proteins were separated in SDS-polyacrylamide gels and transferred to nitrocellulose membranes using standard procedures. Membranes were incubated at 4C overnight with primary antibodies (anti-RAD21; Thermo Fisher PA-5-28344) diluted 1:1000 in 5% milk/TBST. Secondary donkey anti-Rabbit antibody conjugated to HRP (Thermo Fisher) was diluted 1:10,000 in 5% milk/TBST and incubated for 2 hours at room temperature. Immunoblots were developed by applying SuperSignal West Femto Substrate (Thermo Fisher). Immunoblots were visualized using an Odyssey Fc imaging system (LI-COR).

### Generation of clonal cell lines

CTCF-FKPB12^F36V^-mScarlet and Rad21-FKPB12^F36V^-mScarlet cell lines were constructed using CRISPR-Cas9. Previously published targeting guide RNAs and homologous recombination repair templates were used, except that GFP tags were replaced with mScarlet in repair templates ^43^. Vectors were isolated using either CsCl purification or with Genejet Endo-free Plasmid Maxiprep Kit. 15 μg donor repair plasmid and 5 μg sgRNA plasmid were transfected into pro-B cells using Neon transfection system (conditions: 1650 V, 1 pulse, 20 ms). 2-3 days post transfection, mScarlet expressing single cells were sorted into individual wells of 96-well plates. 7-10 days post-sort, individual colonies were screened by PCR to verify proper insertion of FKPB12^36V^-mScarlet.

### SNAP-Tag labeling

To label mut-TetR-SNAP expressing cells, SNAP-Cell 647-SiR was added to cell culture media at a 1:2500 dilution and incubated at 37C for 30 min. After 30 min, cells were spun down and rinsed 3X with 10 mL fresh culturing media. Cells were then incubated in 10 mL fresh culturing media for 4-6 hours to allow unbound SNAP-Cell 647-SiR to diffuse out of the cell. Cells were then spun down and rinsed 1X with 10 ml fresh culturing media.

### Imaging

Cells were plated in FluoroDish, Poly-D-lysine Sterile Culture Dishes (World Precision Instruments) for imaging. Imaging was performed on the Zeiss Elyra 7 Lattice SIM super resolution microscope (100X 1.4 NA oil immersion objective). Cells were supplemented with 5% CO2 while imaging, and the stage and objective were heated to 37C. 1% and 1.5% laser powers were set for 488 and 647 lasers, respectively, with a 40 ms exposure time. Far red and green channels were simultaneously acquired with dual cameras. Z-stacks were acquired with a 0.200 μm spacing between steps. Z-stacks were taken every 4 seconds for a total of 800 seconds. Three fluorescent foci were detected, reflecting the insertion of an array of MUT-TET operator sites on a single allele, as previously reported ^27^.

### Tracking

Positions of fluorescent WT-TetR-eGFP and mut-TetR-SNAP foci were extracted using Imaris microscopy image analysis software. Gaussian filters and background subtraction were applied to help resolve fluorescent puncta. The x, y and z positions of the puncta were determined by applying an Intensity Center of Mass filter. A track duration filter was applied to remove false signals. Tracks for each signal (two DJ labeled loci and one distal V labeled loci) were synchronized using previously published code ^27^. Only tracks in which all three signals were detected for more than 100 time points were included in the analysis.

### Localization Error

Localization error was calculated from the MSD of the apparent motion in fixed cells. Cells were labeled with SNAP-Cell 647-SiR prior to fixation. Cells were first attached to poly-L-lysine coated slides by incubating cells on slides for 30 min at 37C. Slides were then rinsed in PBS and fixed with 4% PFA for 10 min. Fixation was quenched with 0.1M Tris-HCl, pH 7.4 for 10 min and slides were rinsed again with PBS. Cells were stained with GFP Recombinant Rabbit Monoclonal Antibody (catalog# G10362) at a 1:2500 dilution at 4 ^0^C overnight and with Alexa Flour 488 donkey anti-rabbit IgG for 1 hr. Imaging conditions and tracking conditions were the same as for live cell imaging. Localization error was calculated by calculating the MSD of observed genomic motion for the tracked fixed loci and was determined to be ∼0.003 μm^2^ (Figure S7).

### Mean squared displacement

MSD of the observed apparent genomic motion was calculated as the mean squared change in the spatial separation *r* between the intrachromosomal distal V_H_ and D_H_J_H_ regions, MSD = 〈(*r*(*t*) − *r*(*t* + *τ*))^2^〉. Intrachromosomal V_H_ and D_H_J_H_ gene pairs could be unambiguously identified as the pairs in closer spatial proximity. MSD of the true relative genomic motion was determined from the MSD of the observed apparent motion as previously described ^27^, using the localization error determined in this study. Using the same approach, MSDs for interchromosomal D_HJH–_D_H_J_H_ motion were computed (Figure S8). The values of the scaling exponents and diffusion coefficients were extracted from MSD curves using non-linear least squares fit.

### Identification of potential encounters

Genomic encounters were defined as events where the V_H–_D_H_J_H_ distance falls below a threshold set by the interaction range of the chromatin fiber, with correction for localization error. Specifically, potential encounters were identified as events for which the distance between the V_H_ and D_H_J_H_ regions dropped to or below a threshold r*, where r* = (r_int2_ + r_err2_)^1/2^. We used the values for the interaction distance r_int =_ 100 nm and the localization error r_err =_ 0.05 μm.

## ACKNOWLEDGMENTS

We thank Chris Benner for help with designing a diagram depicting the immunoglobulin heavy chain locus. This research was supported by Simons Investigator Award 929737 (to O.K.D), National Science Foundation EAGER Award 2232049 (O.K.D), National Institutes of Health grants R01 AI082850 and 1R21 AI193549 (C.M.), and UC San Diego School of Medicine Microscopy Core Grants P30 NS047101 and S10 OD030505. This study was also supported by a gift from Edwina Riblet in honor of Dr. Roy Riblet (C.M.).

## AUTHOR CONTRIBUTIONS

Conceptualization, O.K.D., M.A and C.M.; experiments, M.A. performed the majority of the experiments; A.P. performed qPCR; investigation, O.K.D., M.A and C.M.; analysis, M.A. and O.K.D; writing, M.A., O.K.D. and C.M.

## DECLARATION OF INTERESTS

The authors declare no competing interests.

**Table S1. Extracted diffusion coefficients and scaling exponents.** a) Generalized diffusion coefficients (D, μm^2^/s^α^) and scaling exponents (α) extracted from MSD of the trajectories measured under each of the experimental conditions over short (4 -80 seconds) and long (100 -400 seconds) times.

b) Diffusion coefficients (D) and scaling exponents (α) extracted from the sub-population of trajectories with large spatial separations (r > 1.25 μm) in RAD21-depleted cells. Reported uncertainties represent fitting errors obtained from the nonlinear least-squares fits of the MSD curves.

**Figure S1. Inhibition of transcriptional elongation in pro-B cells interferes with Igh locus non-coding transcript abundance.**

a) Diagram depicting the folding patterns of the Igh locus fiber in pro-B cells. Bound CTCF motifs are indicated for sense (blue) and anti-sense orientation (brown). Genomic position of the WT-TET (green) and MUT-TET operator sites in the Igh locus are shown. Non-coding transcripts Im and PAIR 4 are indicated. Cartoon was modified from Benner et al. ^24^.

b) ARP, b-actin, Im and PAIR 4 transcript levels are shown for WT-TET;MUT-TET pro-B cells that were cultured in the absence or presence of the transcriptional elongation inhibitor DRB.

**Figure S2. Quantification of transcription-dependent changes in trajectory demixing.**

a) Distribution of the normalized trajectory width, (*q*_90_ − *q*_10_)/median(*r*), for individual V_H–_D_H_J_H_ trajectories under DMSO and DRB conditions, where *r*(*t*) is the distance trajectory and *q*_10_ and *q*_90_ denote its 10th and 90th percentiles. Transcription inhibition significantly increased the normalized trajectory width (permutation test *p* = 0.0046), indicating that individual V_H–_D_H_J_H_ pairs explore a broader range of spatial separations.

b) V_H–_D_H_J_H_ distance trajectories under transcription inhibition (DRB). Gray lines show all trajectories. Red and blue lines indicate trajectories exhibiting the largest increases and decreases, respectively, in spatial separation between the beginning and end of the observation window.

**Figure S3. Individual trajectories in BCR-ABL transformed RAG2-/-pro-B cells carrying the CTCF-FKBP12^F36V^-mScarlet degron.**

Temporal trajectories of V_H_-D_H_J_H_ distances for cells carrying the CTCF-FKBP12^F36V^-mScarlet degron cultured in the presence of DMSO or dTAG-13 as shown in Figures 3e and 3f, but including a single trajectory associated with > 3 μm spatial V_H_-D_H_J_H_ separation when cultured in the presence of DMSO. Trajectories were color-coded based on the mean value of V_H_-D_H_J_H_ distance and plotted as a function of time (s).

**Figure S4. Rapid depletion of RAD21 in pro-B cells carrying a** RAD21-FKBP12F36-mScarlet degron. Western blot analysis to monitor RAD21 expression in BCR-ABL transformed WT-TET; MUT-TET RAG2-deficient pro-B cells and in BCR-ABL transformed RAG2-deficient pro-B cells that carried the RAD21-FKBP12F36-mScarlet degron. Cells were cultured in the absence versus presence of dTAG-13. Molecular weights of wild-type RAD12 and RAD21-FKBP12-mScarlet are indicated.

**Figure S5. Loss of self-similarity in the trajectories with large spatial separations is not caused by small sample size.**

Rescaled velocity correlation functions for groups of 10 randomly sampled trajectories recorded in DMSO-treated BCR-ABL transformed RAD21-FKBP12^F36V^-mScarlet pro-B cells carrying tandem arrays of WT-TET and MUT-TET operator binding sites.

**Figure S6. Paired V_H_ and D_H_J_H_ elements that are located within close spatial proximity are less mobile than paired V_H_ and D_H_J_H_ elements separated by larger distances.**

a-b) Time-and-ensemble averaged mean-square displacement for trajectories associated with mean spatial V_H_-D_H_J_H_ distances greater than 500 nm (dark grey) and less than 500 nm (light grey) for BCR-ABL transformed RAG2-/-pro-B cells that carry the WT-TET and MUT-TET operator binding sites when cultured in the presence of DMSO or DRB.

c-d) Time-and-ensemble averaged mean-square displacement for trajectories associated with mean spatial V_H_-D_H_J_H_ distances greater than 500 nm (dark grey) and less than 500 nm (light grey) for BCR-ABL transformed RAG2-/-CTCF-FKBP12^F36V^-mScarlet pro-B cells that carry the WT-TET and MUT-TET operator binding sites when cultured in the presence of DMSO (left) or dTAG-13 (right).

e-f) Time-and-ensemble averaged mean-square displacement for trajectories associated with mean spatial V_H_-D_H_J_H_ distances greater than 500 nm (dark grey) and less than 500 nm (light grey) for BCR-ABL transformed RAD21-FKBP12^F36V^-mScarlet pro-B cells that carry the WT-TET and MUT-TET operator binding sites when cultured in the presence of DMSO (left) or dTAG-13 (right.

**Figure S7. Mean-square displacement curves associated with V_H_-D_H_J_H_ and D_HJ_H-D_H_J_H_ motion in formaldehyde fixed pro-B cells**

a) Time-and-ensemble averaged MSD curves associated with V_H_-D_H_J_H_ trajectories adopted in formaldehyde-fixed BCR-ABL transformed pro-B cells that carry WT-TET and MUT-TET operator binding sites.

b) Time-and-ensemble averaged MSD curves associated with D_HJH-_D_H_J_H_ trajectories adopted in formaldehyde-fixed BCR-ABL transformed RAG2 -/-pro-B cells that carry WT-TET and MUT-TET operator binding sites.

**Figure S8. Average D_HJH-_D_H_J_H_ interchromosomal motion for pro-B cells following inhibition of transcription and depletion of CTCF or RAD21.**

a) Time-and-ensemble averaged MSD curves of true relative **D_HJH-_D_HJH_** _m_otion for BCR-ABL transformed RAG2-deficient pro-B cells when cultured in the presence of DMSO (dark grey) and DRB (dark purple). MSDs prior to correction are also shown (light shades).

b) Time-and-ensemble averaged MSD curves of true relative D_HJH-_D_H_J_H_ motion under DMSO (dark yellow) and dTAG-13 (light yellow) for BCR-ABL transformed RAG-/-pro-B cells that carry the CTCF-FKBP12^F36V^-mScarlet degron. MSDs prior to correction are also shown (light shades).

c) Time-and-ensemble averaged MSD curves of true relative D_HJH-_D_H_J_H_ motion under DMSO (dark red) and dTAG-13 (light red) conditions for BCR-ABL transformed pro-B cells that carry the RAD21-FKBP12^F36V^-mScarlet degron. MSDs prior to correction are also shown (light shades).

## Notes

### Competing Interest Statement

The authors have declared no competing interest.

