## Supplementary Information for "A hierarchy of spatial confinements governs chromatin dynamics and genomic encounters"

**Table S1** ***Aubrey et al***

**a**

| | | 4 - 80 seconds | 100 - 400 seconds |
|---|---|---|---|
| **DMSO (DRB)** | D | 0.0088 ± 0.0004 | 0.0061 ± 0.0004 |
| | α | 0.28 ± 0.01 | 0.36 ± 0.01 |
| **DRB** | D | 0.0127 ± 0.0006 | 0.0035 ± 0.0002 |
| | α | 0.27 ± 0.01 | 0.53 ± 0.01 |
| **DMSO (CTCF)** | D | 0.0070 ± 0.0004 | 0.0073 ± 0.0003 |
| | α | 0.26 ± 0.02 | 0.26 ± 0.01 |
| **dTAG-13 (CTCF)** | D | 0.0103 ± 0.0004 | 0.0060 ± 0.0006 |
| | α | 0.22 ± 0.01 | 0.34 ± 0.02 |
| **DMSO (RAD21)** | D | 0.0065 ± 0.0002 | 0.0098 ± 0.0006 |
| | α | 0.32 ± 0.01 | 0.23 ± 0.01 |
| **dTAG-13 (RAD21)** | D | 0.0090 ± 0.0010 | 0.0133 ± 0.0013 |
| | α | 0.36 ± 0.03 | 0.27 ± 0.02 |

**b**

| | | 4 to 60 seconds | 100 to 400 seconds |
|---|---|---|---|
| **dTAG-13 (RAD21)** | D | 0.0130 ± 0.0026 | 0.0610 ± 0.0101 |
| r > 1.25 μm | α | 0.54 ± 0.05 | 0.14 ± 0.03 |

### Figure S1 *Aubrey et al*

**a**

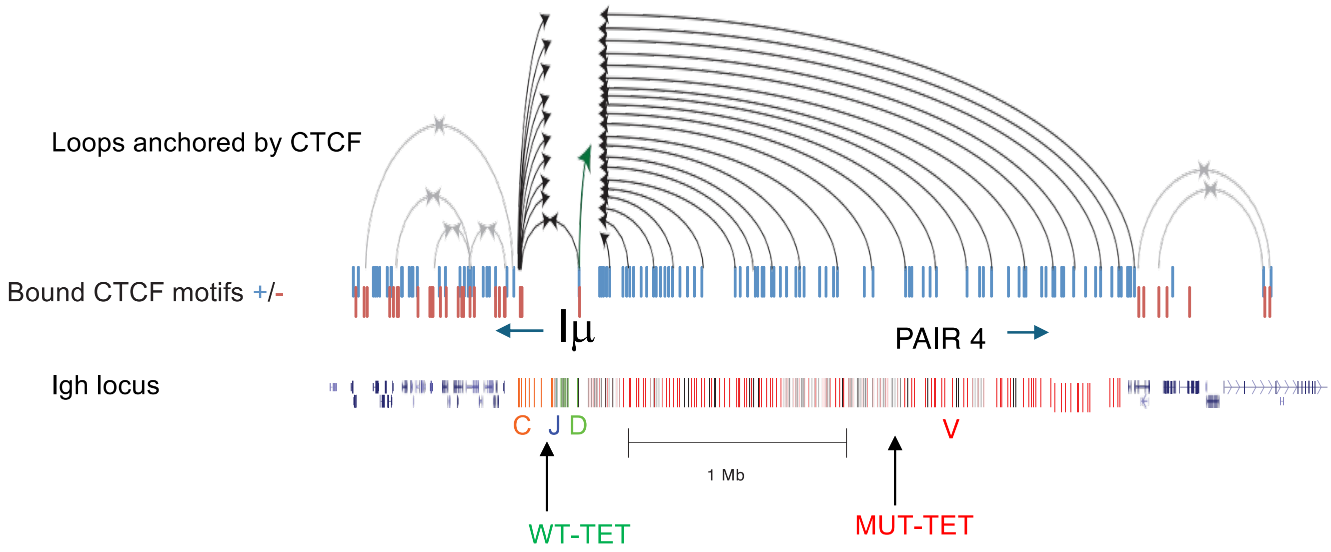


**b**

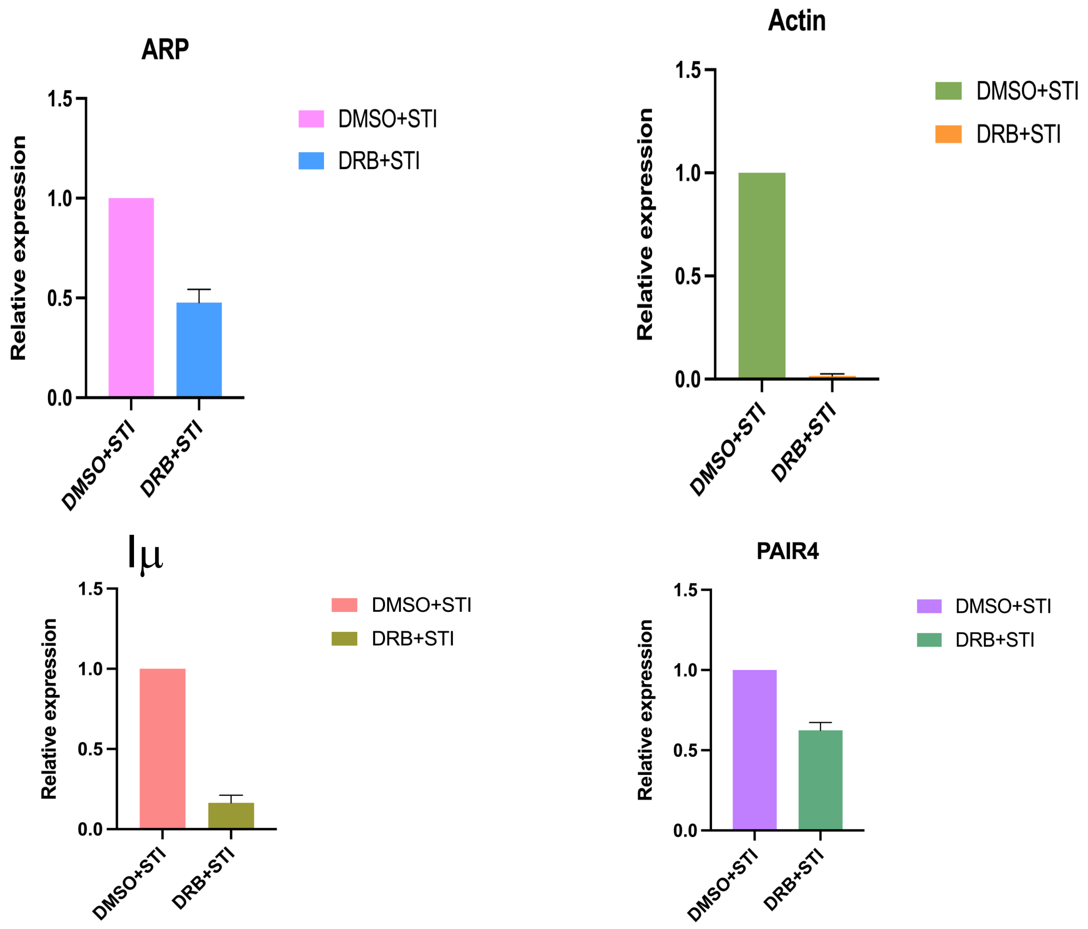

Figure S2 *Aubrey et al*

**a**

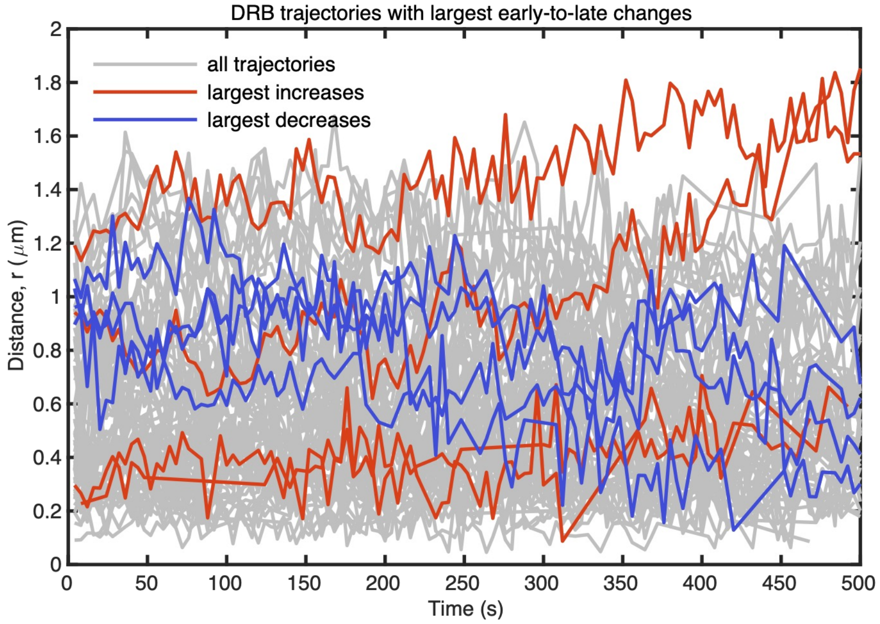


**b**

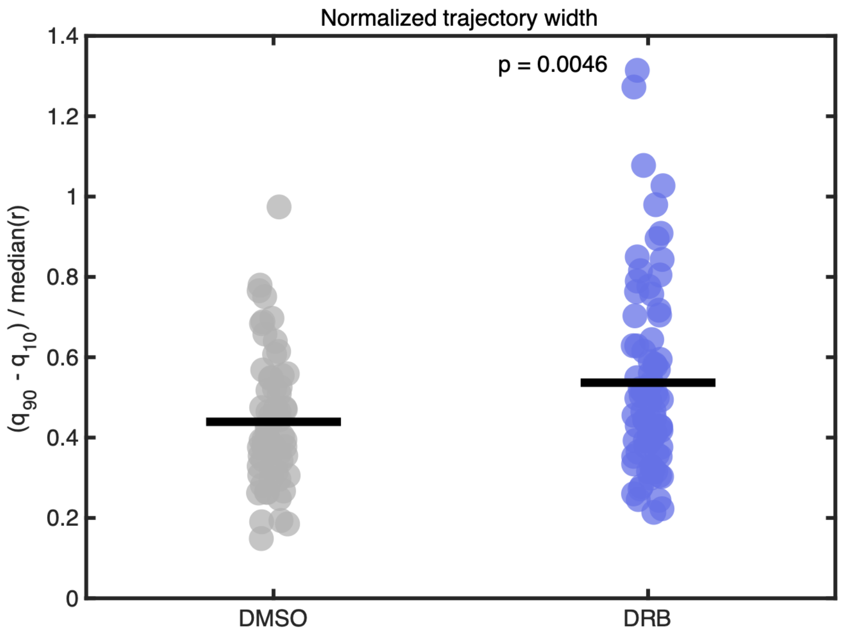

**Figure S3 *Aubrey et al***

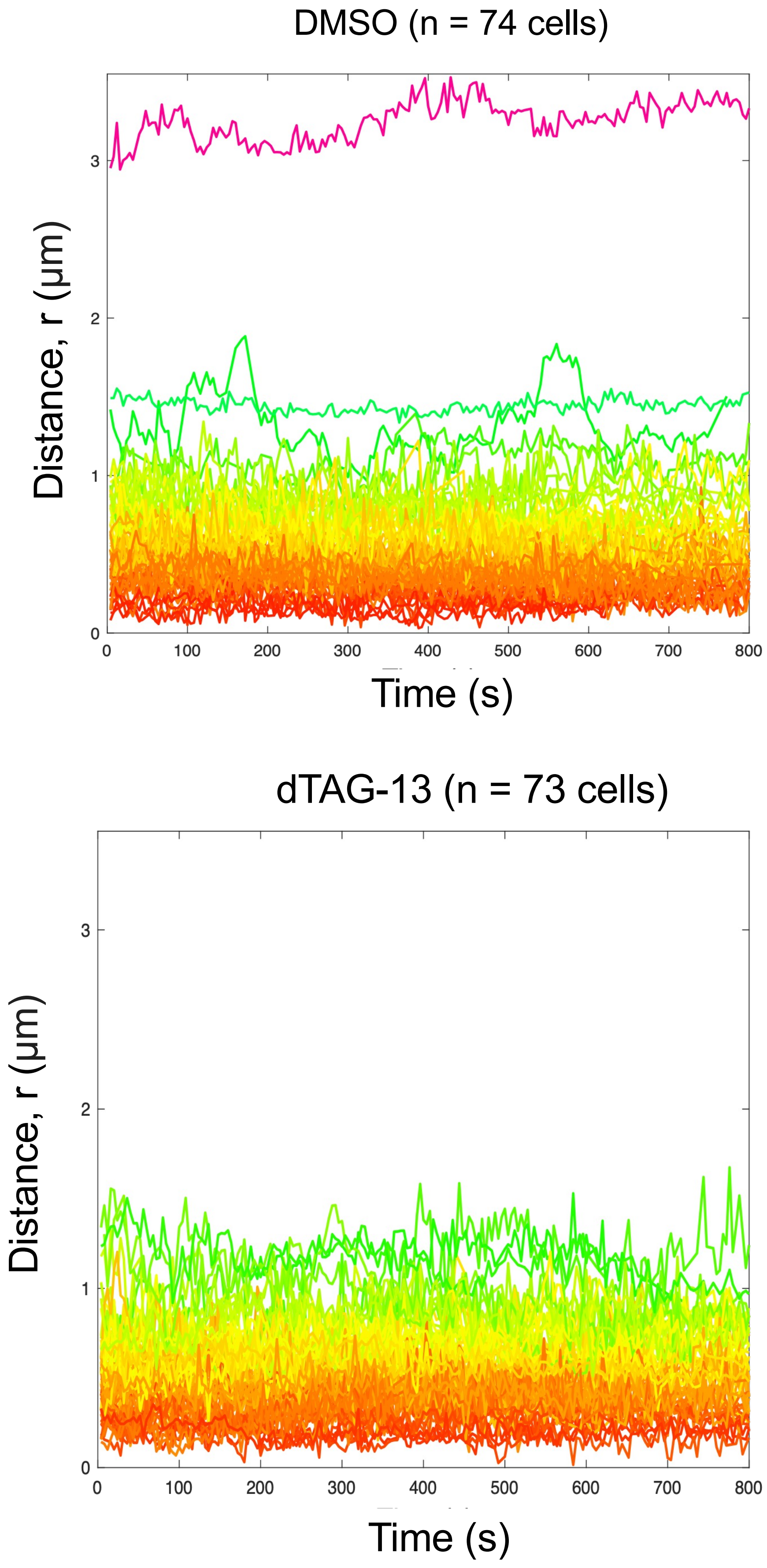

**Figure S4 *Aubrey et al***

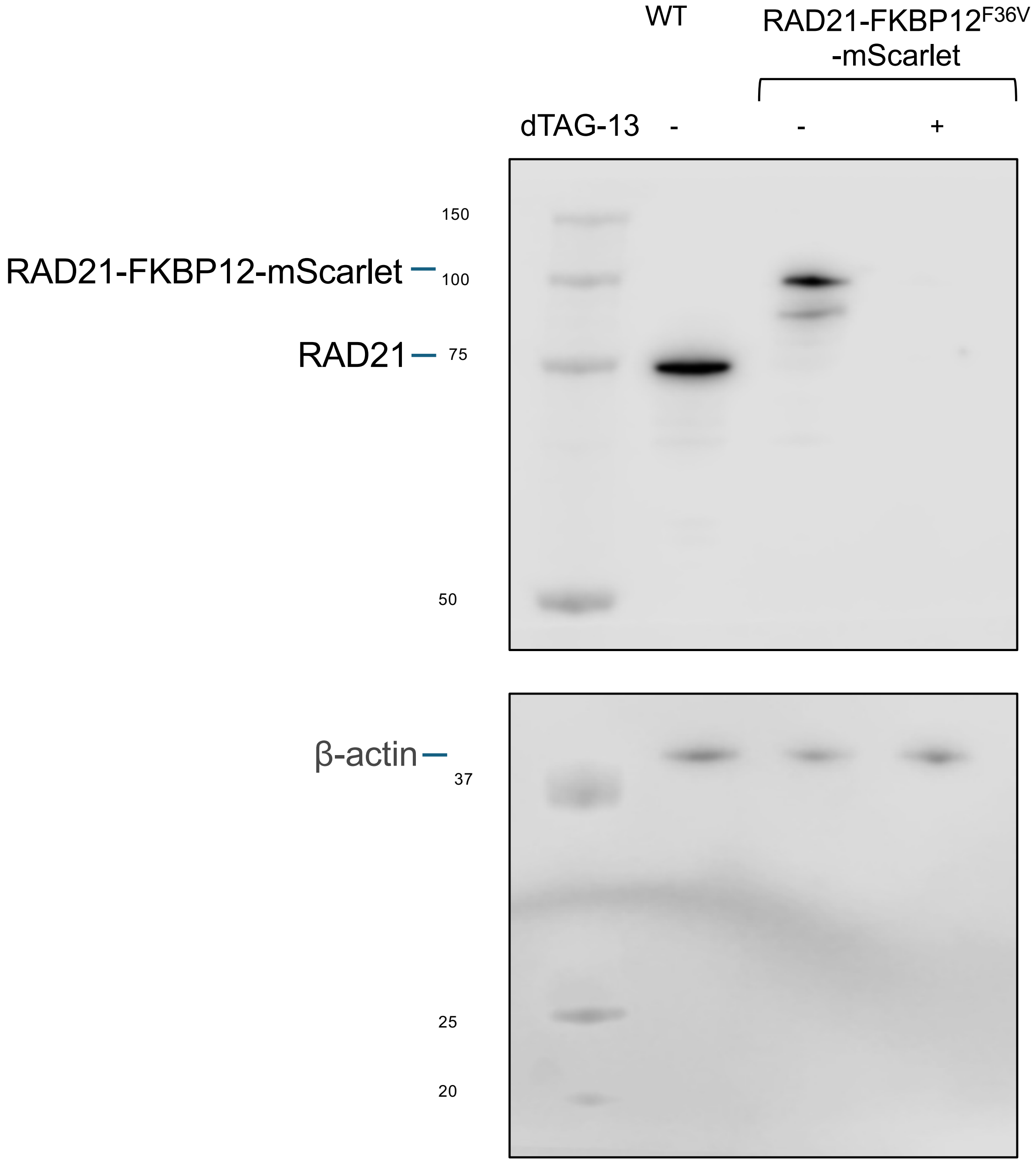

### Figure S5 *Aubrey et al*

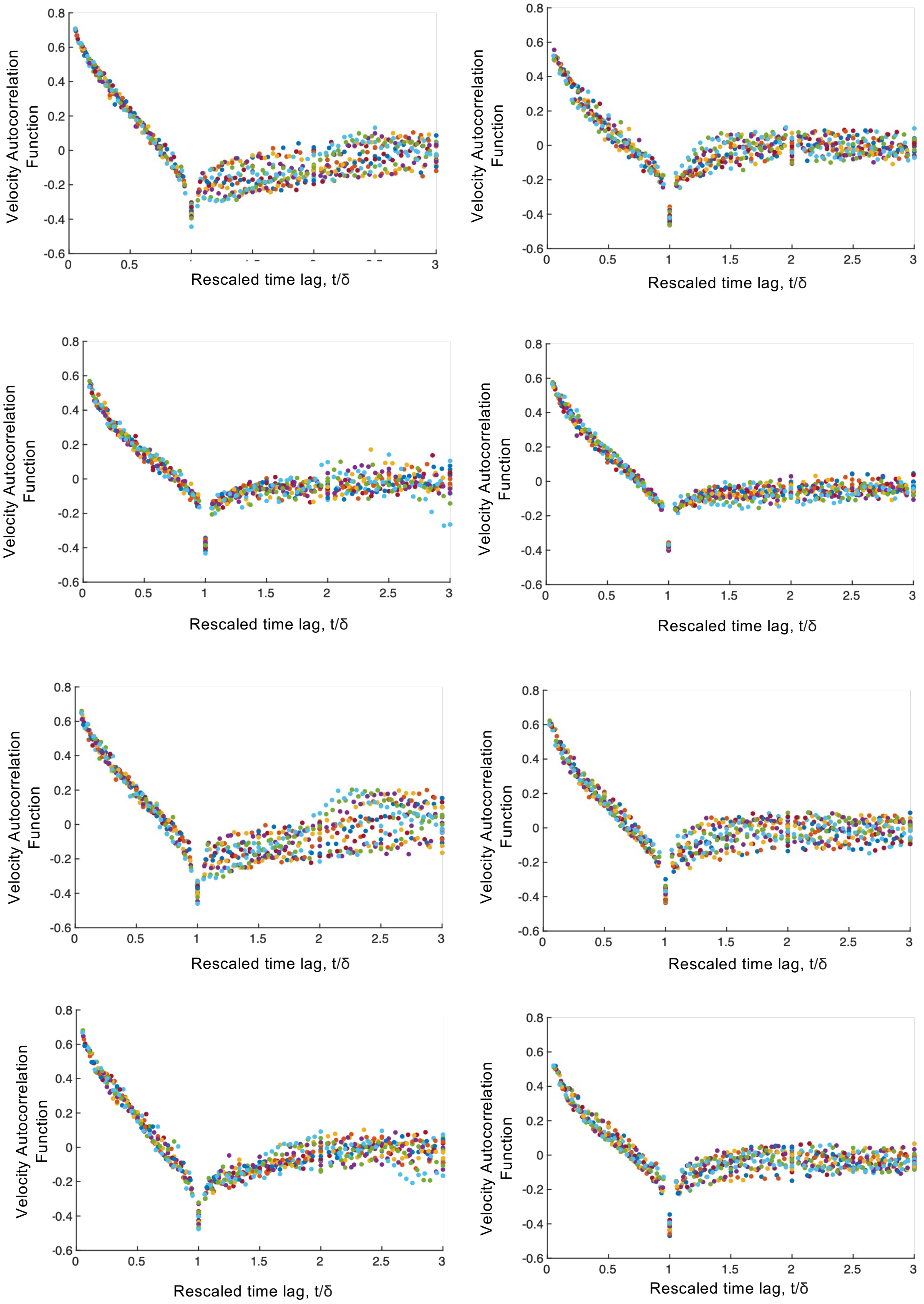

### Figure S6 *Aubrey et al*

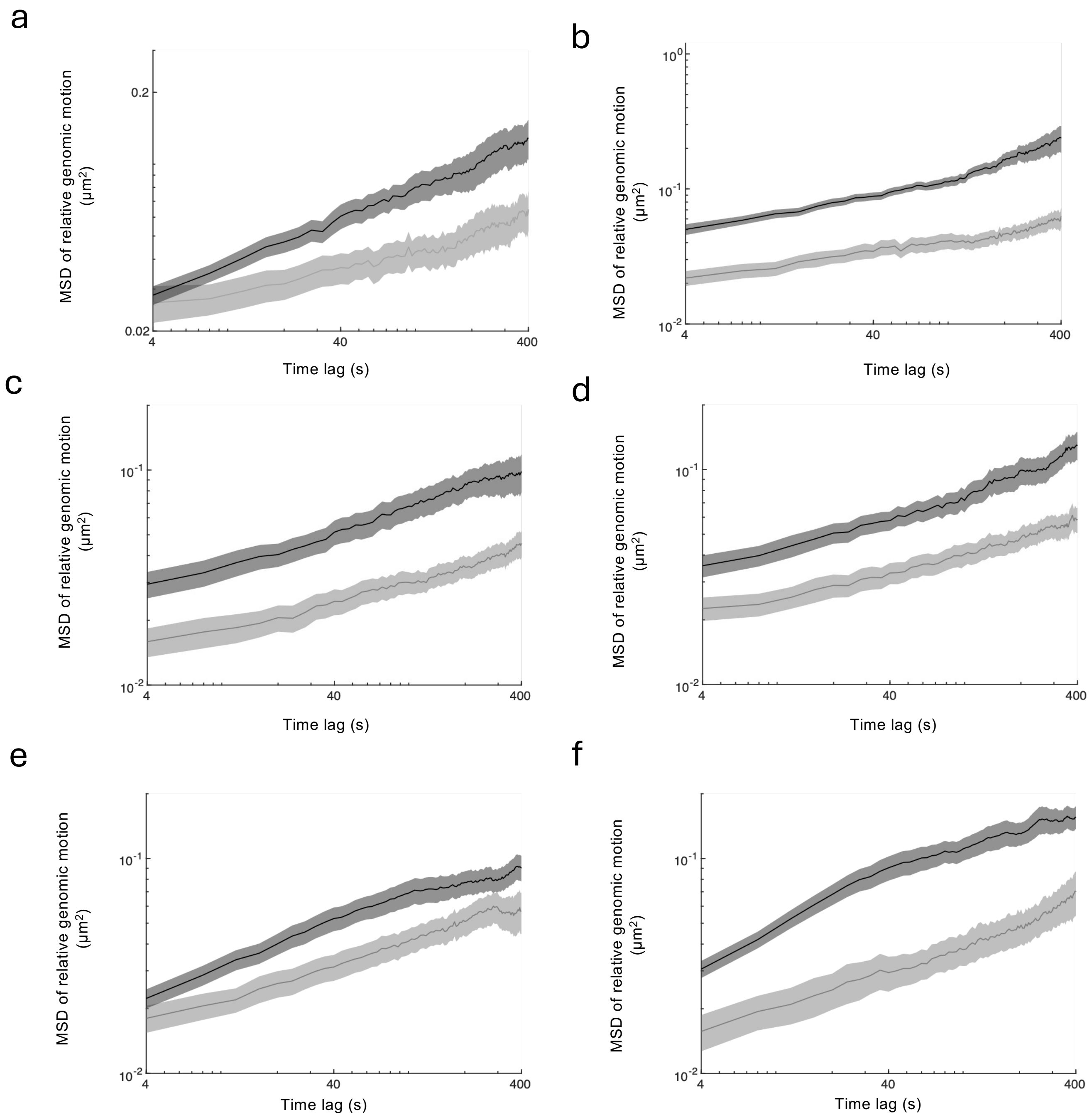

**Figure S7 *Aubrey* *et al***

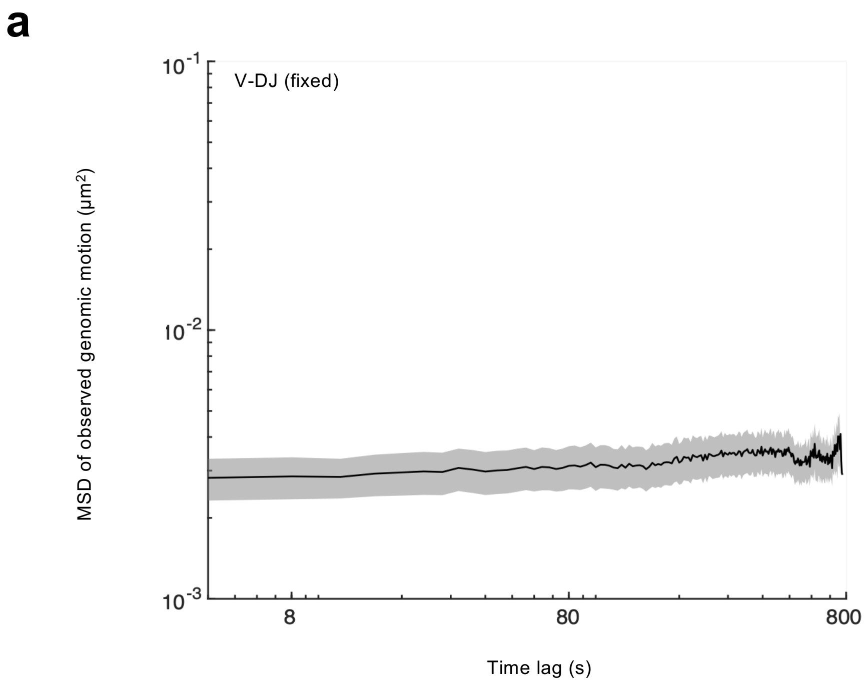


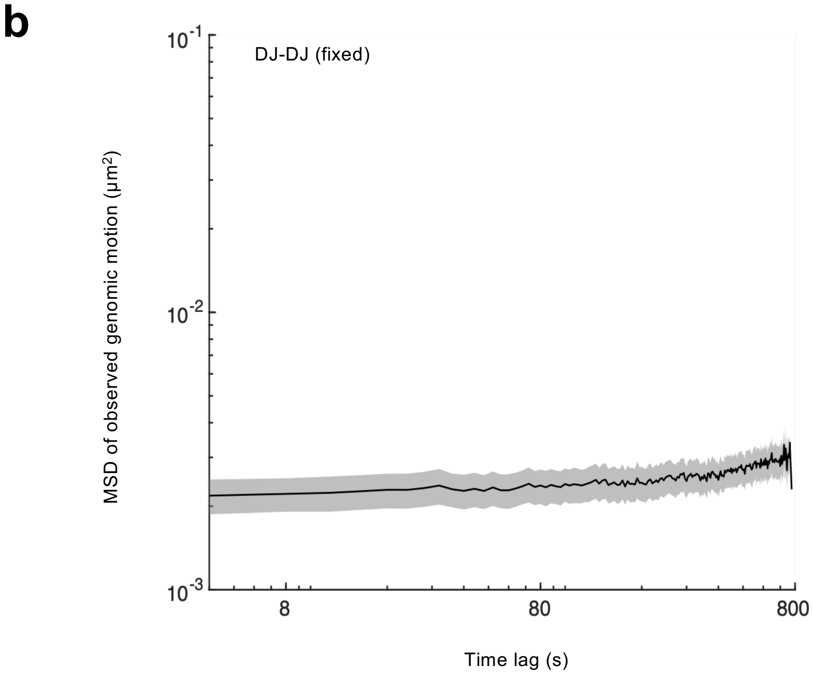

**Figure S8** ***Aubrey et al***

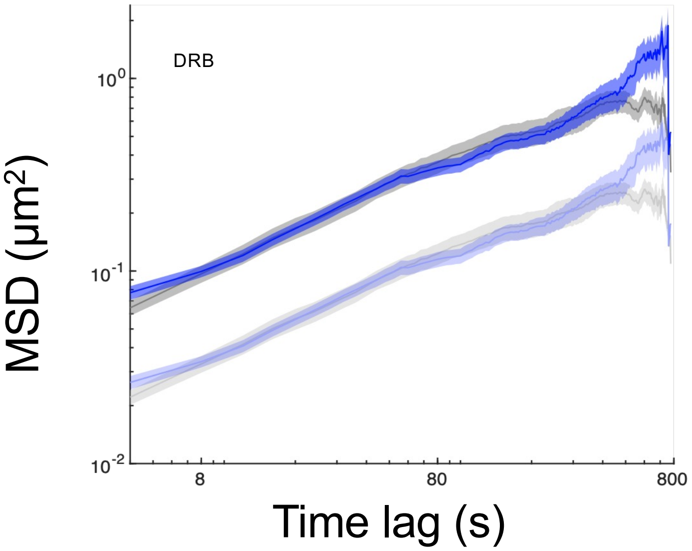


b

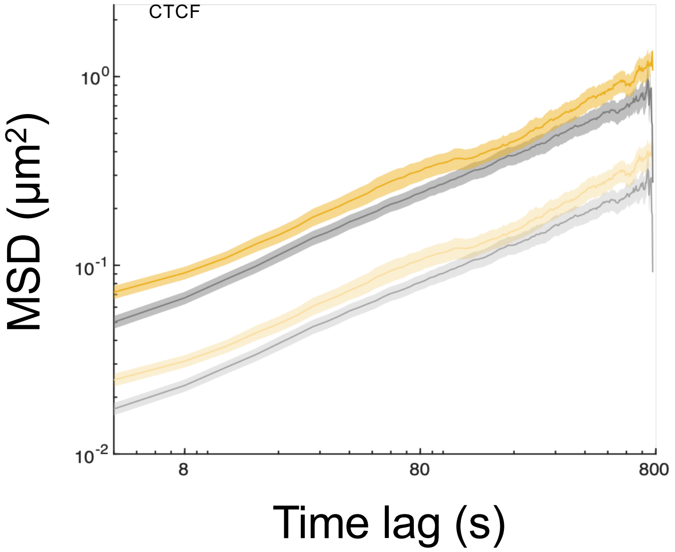


c

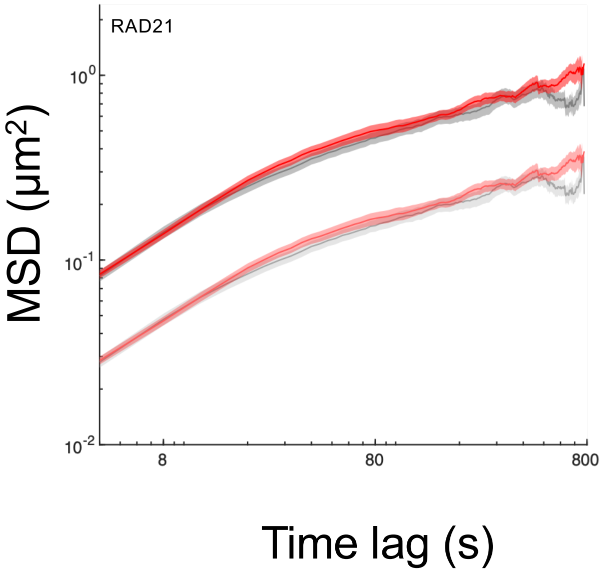
